# MARBL: A Live-Cell Method for Profiling Bioenergetic Heterogeneity by Noncanonical Methionine Labeling of the Cell Surface Proteome

**DOI:** 10.64898/2026.06.01.729353

**Authors:** Lillian R. Delacruz, Ming Ye, Kendra A. Libby, Keran Han, Chad J. Munger, Yunping Qiu, Irwin J. Kurland, Alison E. Ringel

## Abstract

Heterogeneity is a hallmark of biological systems, where cell-to-cell variability supports adaptation to changing environments, but also enables maladaptive states such as drug resistance. Many sources of non-genetic variation, particularly bioenergetics and metabolism, remain difficult to measure in living cells and connect to functional outcomes. Here, we introduce MARBL (Methionine Analogues for Ratiometric Bioenergetics in Live cells), a method that encodes translationally-coupled energetic responses to metabolic stress as an internally normalized signal within the surface proteome of living cells. Applying MARBL to primary immune cells reveals that differences in baseline translational activity can underlie apparent metabolic vulnerabilities, underscoring the importance of ratiometric measurements. We demonstrate that MARBL can enrich pathogenic from non-pathogenic TH17 cells based on resilience to bioenergetic stress, which functionally distinguishes cells that produce IFNγ upon restimulation. Overall, MARBL offers a versatile platform to profile metabolic resilience in living cells and link bioenergetic state to cellular function.

## Introduction

Cells must generate chemical energy to sustain a wide array of biological processes necessary for growth, proliferation, and survival. To do this, cells take up fuels from the local environment and oxidize them through metabolic pathways that maintain high-energy ATP pools. As virtually every cellular process requires chemical energy, cells adjust the activity of energy-producing pathways based on functional demands and fuel availability. Accordingly, bioenergetic programs can be highly dynamic. For example, cells remodel the expression of metabolic machinery in response to external factors like nutrient availability, as well as cell state transitions such as malignant transformation or immune activation (1–7). Experimental tools that assess bioenergetic state and fuel utilization have provided key insights into the reciprocal regulation between metabolic programs and cellular behaviors (3,7–13). However, higher resolution methods that can functionally assess energetic heterogeneity are still needed.

While single-cell genomic, transcriptomic, and epigenomic technologies have yielded remarkable insights into molecular heterogeneity among individual cells, investigating metabolism approaching single-cell resolution has proven more challenging. Current tools that directly quantify ATP levels, such as mass spectrometry, require large quantities of purified cell input and return bulk measurements averaged over all cells in the sample. Similarly, classical bioenergetic assays, where substrates and/or inhibitors are given to bulk populations of cells to infer metabolic state, also provide population-level readouts. Seahorse metabolic flux assays are widely used for this purpose, measuring how extracellular acidification (ECAR) and oxygen consumption rate (OCR), which serve as proxies for glycolytic activity and oxidative phosphorylation, respectively, change upon treating cells with a series of inhibitors. However, population-averaged Seahorse measurements are not easily integrated with downstream functional analyses that require live cells. The recent development of SCENITH is a major advance that overcomes some of these technical barriers for single-cell energetic profiling (11). SCENITH infers bioenergetic state by quantifying puromycin incorporation into nascent polypeptides. This is possible since protein synthesis is one of the most energy-intensive processes in living cells (14). Since puromycin incorporation is measured by flow cytometry, SCENITH enables the analysis of heterogeneous cell populations that can be distinguished by lineage markers. However, cells must be fixed and permeabilized to detect puromycin, which precludes downstream functional assays. Also, this technique does not account for variation in baseline translation rates, a factor that can affect the interpretation of translation changes during metabolic inhibition. Thus, there is still an unmet need for workflows that can define functional consequences of metabolic heterogeneity, as well as to measure these features in rare cell populations.

Here, we have developed a flexible and modular toolkit to monitor cellular energetic state that accounts for variation in basal translation, while also preserving cell viability for downstream functional analyses. Building on the use of protein translation rate as a responsive proxy for bioenergetic state (11,14,15), we have created MARBL (Methionine Analogues for Ratiometric Bioenergetics in Live cells), a workflow to encode normalized translational activity into the surface proteome of live cells. MARBL relies on the incorporation of two clickable methionine analogues (CMAs) with distinct chemical reactivities, enabling paired measurements of baseline and metabolically coupled translation within the same cell (16–23). We extensively optimized copper-catalyzed click chemistry to achieve high signal-to-noise with minimal toxicity, enabling robust fluorescent labeling of surface-exposed CMAs on live cells for detection by flow cytometry. Sequential labeling with two CMAs, where the first captures baseline translation rate and the second measures translation under metabolic perturbation, yields a resilience index that quantifies per-cell ratiometric analysis of bioenergetic resilience under stress. We have applied MARBL to deconvolve metabolic dependencies among mixed cell populations within immortalized mammalian cell lines, primary mouse immune cells, and primary human immune cells. As a major advance, we demonstrate that cells can be sorted and re-cultured based on MARBL signal prior to performing downstream functional assays. In addition to providing a normalized measure of bioenergetic state, this approach can be readily integrated with other live-cell assays, adding metabolic heterogeneity as a new dimension for understanding cell state and function. In summary, MARBL sequentially labels the surface proteome of live cells with two chemically distinct CMAs in sequence, enabling per-cell ratiometric analysis, live-cell sorting, and downstream functional assays.

## Results

### Optimizing the use of clickable methionine analogues to monitor translation in the surface proteome of living cells

To enable live-cell analysis of metabolically coupled translation, we developed a method for measuring nascent protein synthesis of the surface proteome using CMAs. As labeling requires surface-accessible methionine residues, we calculated the abundance of methionine in the extracellular proteome of humans and mice. Methionine accounts for ∼2% of residues in eukaryotic proteins (24) and occurs at similar frequencies within the extracellular versus intracellular domains of proteins that localize to the plasma membrane (Fig. 1A). This methionine abundance is comparable to its ∼2.3% frequency of all amino acids in the entire human proteome (25,26).

**Figure 1:**
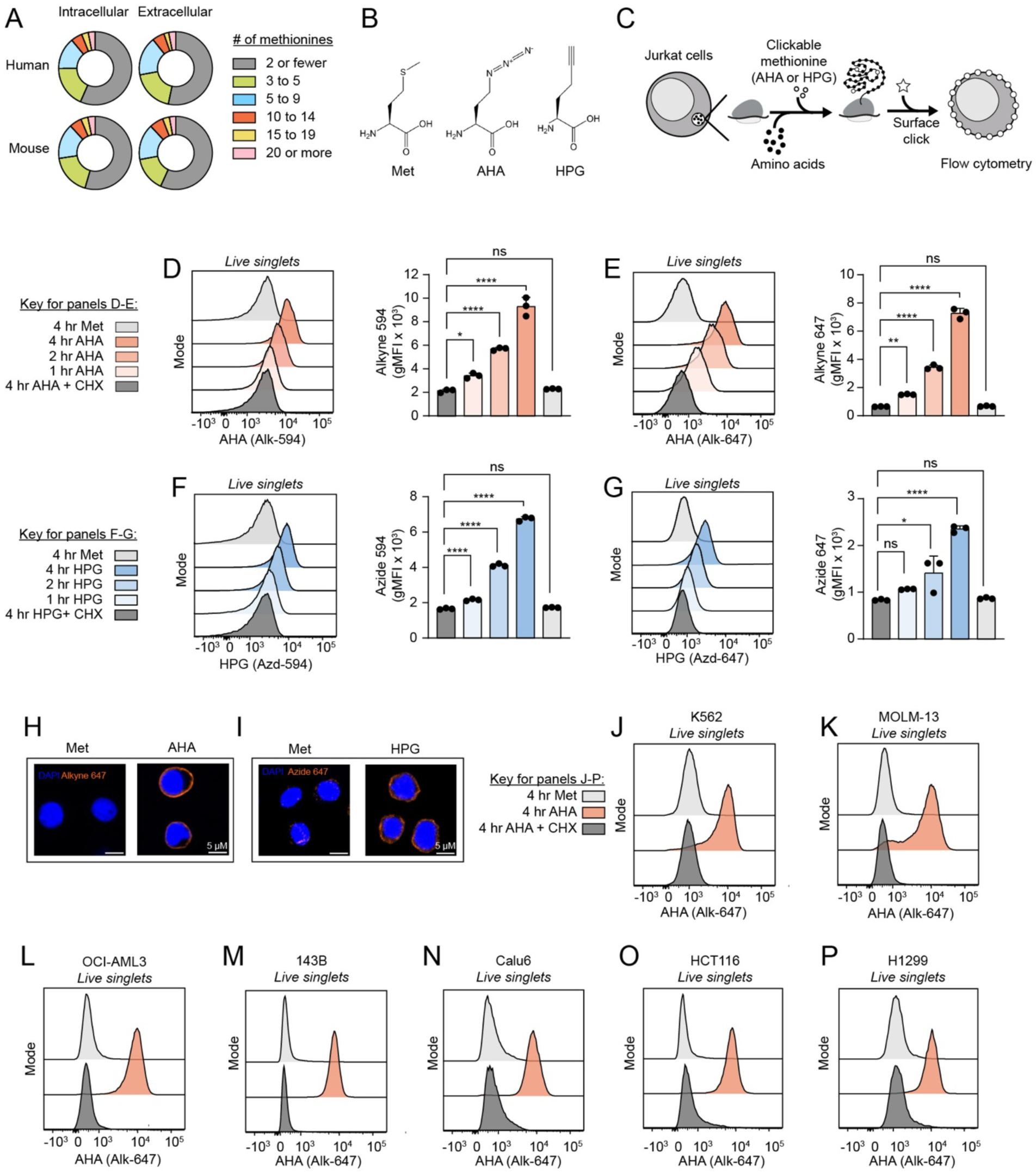
Optimization of clickable methionine analogues to monitor translation in living cells. (A) Frequency of methionine residues in intracellularly or extracellularly localized protein domains within the human or mouse proteins with annotated plasma membrane localization. (B) Chemical structures for methionine and its clickable analogues, AHA and HPG. (C) Schematic depicting experimental workflow. (D and E) Representative flow cytometric histograms and corresponding gMFIs of extracellular Alk-594 signal (D) or Alk-647 signal (E) after 1, 2, or 4 hours of AHA incorporation. (F and G) Representative flow cytometric histograms and corresponding gMFIs of extracellular Azd-594 signal (F) or Azd-647 signal (G) after 1, 2, or 4 hours of HPG incorporation. (H) Representative confocal images of Alk-647 signal, showing surface localization in Jurkat cells after 24 hours of Met or AHA incorporation. (I) Representative confocal images of extracellular Azd-647 signal, showing surface localization in Jurkat cells after 24 hours of Met or HPG incorporation. (J-P) Representative flow cytometric histograms of extracellular Alk-647 signal on K562 (J), MOLM-13 (K), OCI-AML3 (L), 143B (M), Calu6 (N), HCT116 (O), and H1299 (P) cell lines after 4 hours of AHA incorporation. Abbreviations: AHA = Azidohomoalanine, HPG = Homopropargylglycine, Met = Methionine, CHX = Cycloheximide, gMFI = Geometric mean fluorescence intensity Statistical significance was assessed by one-way ANOVA (D-G) followed by Tukey’s multiple comparisons test. Graphs display mean ± SD (D-G). (ns, p >0.05; *, p ≤ 0.05; **, p ≤ 0.01; ***, p ≤ 0.001; or ****, p ≤ 0.0001).

Having established that the mammalian proteome contains methionine residues within extracellular domains, we sought to develop a labeling strategy with chemically reactive CMAs that could be detected on the surface of live cells. We swapped methionine in cell culture medium with two commercially available CMAs with distinct chemical reactivity. Azidohomoalanine (AHA) bears an azide modification while homopropargylglycine (HPG) bears an alkyne modification (Fig. 1B). These analogues are incorporated into nascent proteins by endogenous tRNA(Met) machinery in methionine-free media (27). To detect incorporation into the surface proteome, Jurkat cells were incubated with either AHA or HPG, reacted with chemically compatible fluorophores bearing alkyne or azide moieties using copper-catalyzed azide-alkyne cycloaddition (CuACC) click chemistry (28), and then detected by flow cytometry (Fig. 1C). Upon testing fluorophores with a range of excitation/emission properties, we found that Alk-488, Alk-594, and Alk-647 dyes displayed the highest signal and lowest background staining (Fig. S1A). Hereafter, we use 594 and 647 dyes, but 488 works equivalently well.

Unlike conventional bioorthogonal non-canonical amino acid tagging methods that label the total proteome in cells that have been fixed and permeabilized, the use of intact cells in our workflow restricts labeling to the surface proteome (16,17,29). As targeting surface-exposed methionine residues reduces the total number of reactive moieties available for labeling compared to the entire proteome, we needed to optimize labeling conditions that maximize signal while minimizing background and maintaining viability. After varying incubation times and clicking conditions, we determined that 100 µM of AHA or HPG led to baseline-resolved surface signal in Jurkat cells with minimal impacts on cell viability or growth for up to 24 hours of incubation (Figs. S1B-S1E). To identify the impact of methionine in fetal bovine serum (FBS) on analogue incorporation, Jurkat cells were incubated with AHA or HPG in media supplemented with complete or dialyzed FBS. We observed significantly higher CMA signal in dialyzed serum conditions, indicating that the presence of methionine in complete serum must be avoided (Figs. S1F, S1G). The CuAAC reaction also requires copper(II) ion as a catalyst, which is toxic to cells at the micromolar concentrations needed for CMA labeling (30,31). We tested copper-free alternatives, including dibenzocyclooctyne (DBCO) conjugated fluorophores for AHA labeling, however, the background signal was too high for reliable detection (data not shown). Instead, we pre-complexed Cu^2+^ with a chelating ligand (BTTAA) to shield cells from copper-induced oxidative stress and preserve viability (30). After testing a range of copper concentrations and ligand molar ratios, we found that Jurkat cells clicked with 25 μM CuSO_4_ for AHA and 50 μM CuSO_4_ for HPG, with five-fold excess BTTAA ligand, significantly improved viability (Figs. S1H, S1I). This improvement in cell survival was accompanied by a moderate reduction in Alk-647 signal (on AHA-labeled cells) and enhanced Azd-594 signal (on HPG-labeled cells) (Figs. S1H, S1I). Using these conditions, both AHA and HPG accumulated in a time-dependent manner on the cell surface (Figs. 1D-1G). Addition of cycloheximide (CHX), a compound that inhibits global translation, blocked incorporation of both CMAs (Fig. 1D-J). AHA could be detected equally well when clicked with either Alk-594 or Alk-647 fluorophores (Figs. 1D-E). However, clicking with Azd-594 fluorophore was superior for HPG detection (Figs. 1F-G). For subsequent experiments, we decided to click AHA with alkyne-linked Alk-647 dye and HPG with azide-linked Azd-594 dye. After verifying that this optimized workflow restricts labeling to the surface proteome by microscopy (Fig. 1H-I), we applied it to a variety of suspension (K562, MOLM-13, and OCI-AML3) and adherent (143B, Calu6, HCT116, and H1299) human cell lines, where we detected robust AHA incorporation (Figs. S1J, 1J-1P). Thus, we have optimized an end-to-end workflow to monitor surface translation and preserve viability with CMAs in a variety of cell lines.

### Validation of surface translation as a readout of cellular energetics

Translation is the single largest energy-consuming process in cells, requiring both ATP and GTP to incorporate each amino acid into growing polypeptides (14,32). SCENITH was the first experimental technique to recognize that the global rate of protein synthesis can be used as a proxy for cellular bioenergetic state (11). Therefore, we set out to identify if extracellular incorporation of CMAs likewise reflected cellular energetics. Jurkat cells were incubated with AHA for two hours and metabolic inhibitors that impair cellular bioenergetics were added during the last hour (Fig. 2A). We titrated 2-Deoxy-D-Glucose (2DG) and oligomycin A (Omy) to inhibit glucose metabolism and oxidative phosphorylation (OXPHOS), respectively (Fig. 2B). The resulting cells were processed in parallel for analysis by flow cytometry or quantitative LC-MS/MS. Addition of 2DG and Omy led to a concentration-dependent reduction in AHA incorporation into the surface proteome by flow cytometry (Fig. 2C). The adenylate energy charge (AEC), a ratio that quantifies relative proportions of high-energy nucleotide di- and tri-phosphates compared to the total nucleotide pool, also decreased in a dose-responsive manner with 2DG and Omy treatment and displayed a significant positive correlation with the extracellular click signal (Fig. 2D). In addition, the AHA click signal significantly correlated with isotopic labeling of newly synthesized ATP by H_2_^18^O water labeling (Fig. 2E). These data reveal that surface incorporation of CMAs linearly tracks with energetic adaptations observed with quantitative metabolomics and flux measurements of ATP turnover, both of which are gold standard methods to assay ATP pools.

**Figure 2:**
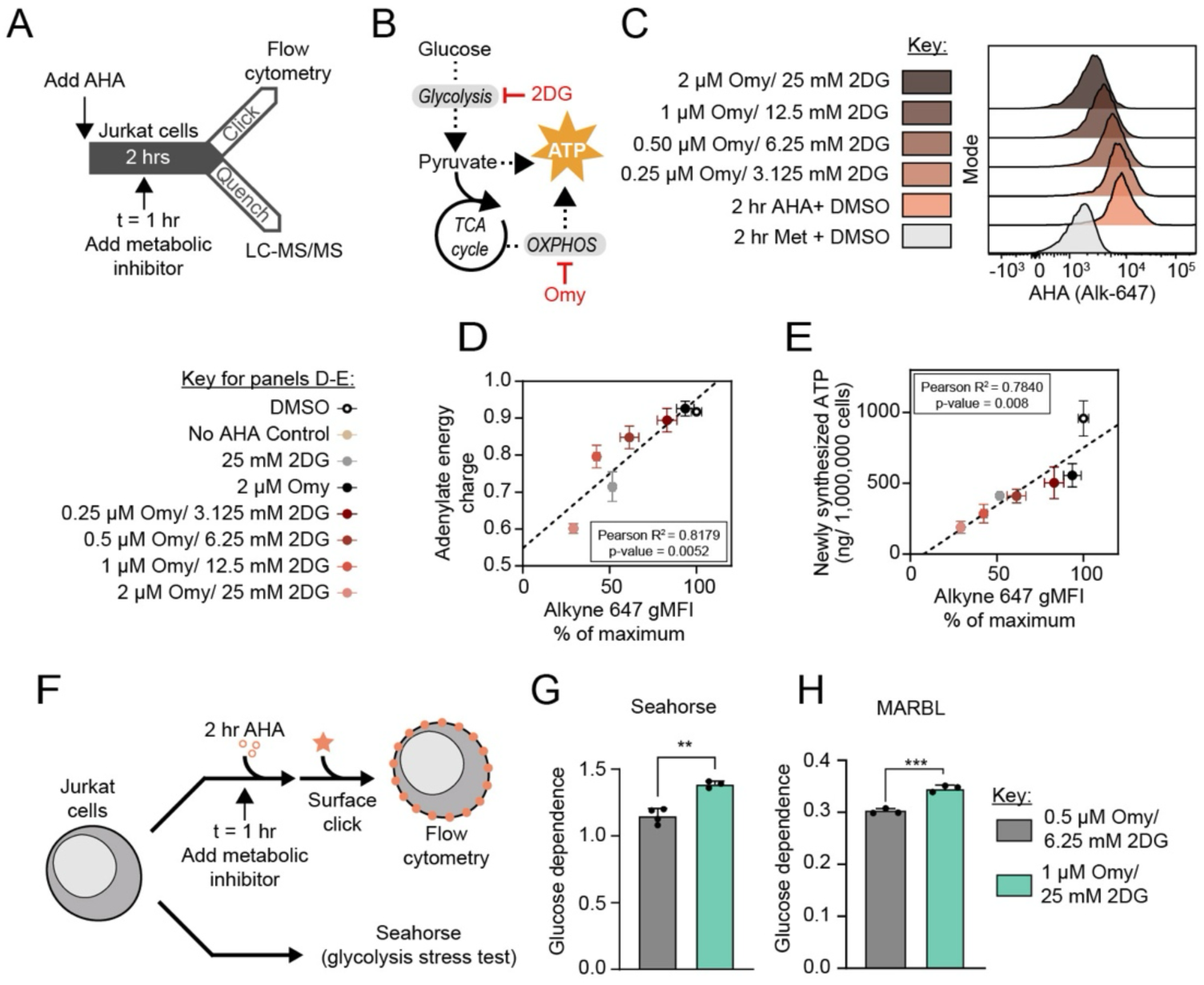
Validation of surface translation rate as a readout of cellular energetics. (A) Schematic depicting experimental workflow. (B) Schematic of ATP-producing pathways and corresponding metabolic inhibitors. (C) Flow cytometric histograms of extracellular Alk-647 signal on Jurkat cells incubated in AHA with varying concentrations of 2DG and Omy. (D) Correlation between changes in adenylate energy charge versus normalized Alk-647 Alkyne flow cytometric signal. (Pearson R^2^ = 0.8179, p = 0.0052). (E) Correlation between changes in newly synthesized ATP versus normalized Alk-647 Alkyne flow cytometric signal (Pearson R^2^ = 0.7840, p = 0.008). (F) Schematic depicting experimental workflow. (G-H) Seahorse glycolysis stress test (G) or MARBL-processing (H) on Jurkat cells using varying concentrations of 2DG and Omy.

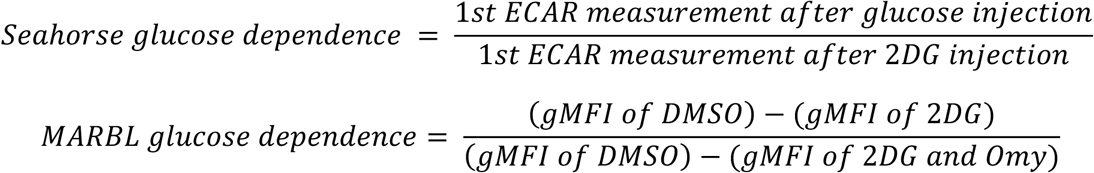 Abbreviations: AHA = Azidohomoalanine, HPG = Homopropargylglycine, Met = Methionine, CHX = Cycloheximide, gMFI = geometric mean fluorescence intensity, 2DG = 2-Deoxy-D-Glucose, Omy = Oligomycin A The Pearson correlation coefficient was used to assess correlation and significance (D-E). Statistical significance was assessed by two-tailed Student’s unpaired t-test (G-H). Graphs display mean ± SD (D-E, G-H). (ns, p >0.05; *, p ≤ 0.05; **, p ≤ 0.01; ***, p ≤ 0.001; or ****, p ≤ 0.0001).

In addition to its role in protein synthesis, methionine participates in one-carbon metabolism, lipid synthesis, and redox regulation. We therefore examined the effect of replacing methionine with AHA or HPG on the polar metabolome at steady state. We performed targeted LC-MS/MS on Jurkat cells grown in CMA-supplemented media across an eight-hour time course. We detected 434 metabolites and observed a significant reduction in methionine as well as metabolites synthesized directly from methionine (methylthioadenosine and S-adenosyl methionine) upon either methionine depletion or CMA treatment across all time points (Fig. S2A). Consistent with the flow cytometry (Figs. 1D-G), cells successfully take up AHA and HPG in as little as two hours of incubation (Fig. S2B). The significant differences common among AHA, HPG, and methionine-free conditions are likely caused by the absence of canonical methionine in media, whereas AHA- or HPG-specific changes may arise from properties of the CMA (Fig. S2C-S2CJ). Most metabolites are unchanged overall, although there are some AHA- and HPG-specific differences (Fig. 2K-M). In sum, we demonstrate that short (hours) incubations in CMAs alter methionine metabolism, with minimal other effects on the basal metabolome in Jurkat cells.

To further benchmark this technique, we compared bioenergetic readouts obtained by surface clicking with outputs from a Seahorse glycolytic stress test (Fig. 2F). Jurkat cells are highly reliant on glucose utilization to fuel energy production via glycolysis (Fig. S2N). In addition, ECAR is insensitive to OXPHOS inhibition with Omy, suggesting that Jurkat cells are unable to raise glycolytic activity any further (Fig. S2N). Therefore, we evaluated glucose dependence by calculating the difference in the ECAR measurement following glucose addition and after 2DG injection in the presence of Omy. Increasing 2DG raised glucose dependence to a similar extent by Seahorse (Fig. 2G) compared to MARBL (Fig. 2H), providing further support that our technique accurately reflects cellular energetics and quantifies it in a dose-dependent manner like a Seahorse assay.

### Development of dual-labeling MARBL workflow to measure cellular energetics in single cells

We next sought to use CMA surface labeling as an internally normalized assay for assessing cellular energetic sensitivity. This requires distinguishing translation variation driven by non-metabolic sources from effects coupled to bioenergetics. To achieve this, we leveraged the distinct chemical reactivities of AHA (which reacts with an alkyne) versus HPG (which reacts with an azide) to measure translation rates under two conditions in the same cell (Fig. 3A). Cells are first incubated in HPG-supplemented media in the absence of drug to capture translation rates at baseline, which provides an internal control for normalization. Next, the same cells are swapped into AHA-supplemented media in the presence of one or more metabolic inhibitors to assess the effect of energetic stress on translation. Finally, surface CMA incorporation is detected via sequential clicking before analysis by flow cytometry. We applied this dual-labeling scheme to Jurkat cells and detected both HPG and AHA incorporation on the cell surface (Fig. 3B). We did not observe any cross-reactivity between clickable fluorophores and largely maintained cell viability (only ∼5% reduction) between the singly versus dually clicked cells (Fig. 3C). HPG signal remained steady while responsiveness to metabolic or translation perturbations was only observed in the AHA signal (Fig. 3D). Providing CHX during AHA inhibition completely blocked the AHA signal, whereas bioenergetic inhibition by treatment with 2DG and Omy resulted in an intermediate reduction of the AHA signal (Fig. 3D). CHX treatment was more toxic compared to 2DG and Omy (Fig. 3E). Next, we tested whether altering either the sequence of CMA addition or the order of surface-clicking reactions impacts performance of the dual-label workflow. The swapped labeling scheme also produced baseline-resolved and metabolically coupled signals (Fig. S3A-S3C). However, AHA displayed superior incorporation, signal, and dynamic range in measuring the translational response to energetic perturbation (Fig. 3B-D). Swapping the sequence of clicked fluorophores while maintaining the HPG followed by AHA labeling sequence results in similar baseline and metabolically coupled signals (Fig. S3D-S3F). Thus, the sequence of the CMAs can impact the dynamic range in a dual-incubation workflow.

**Figure 3:**
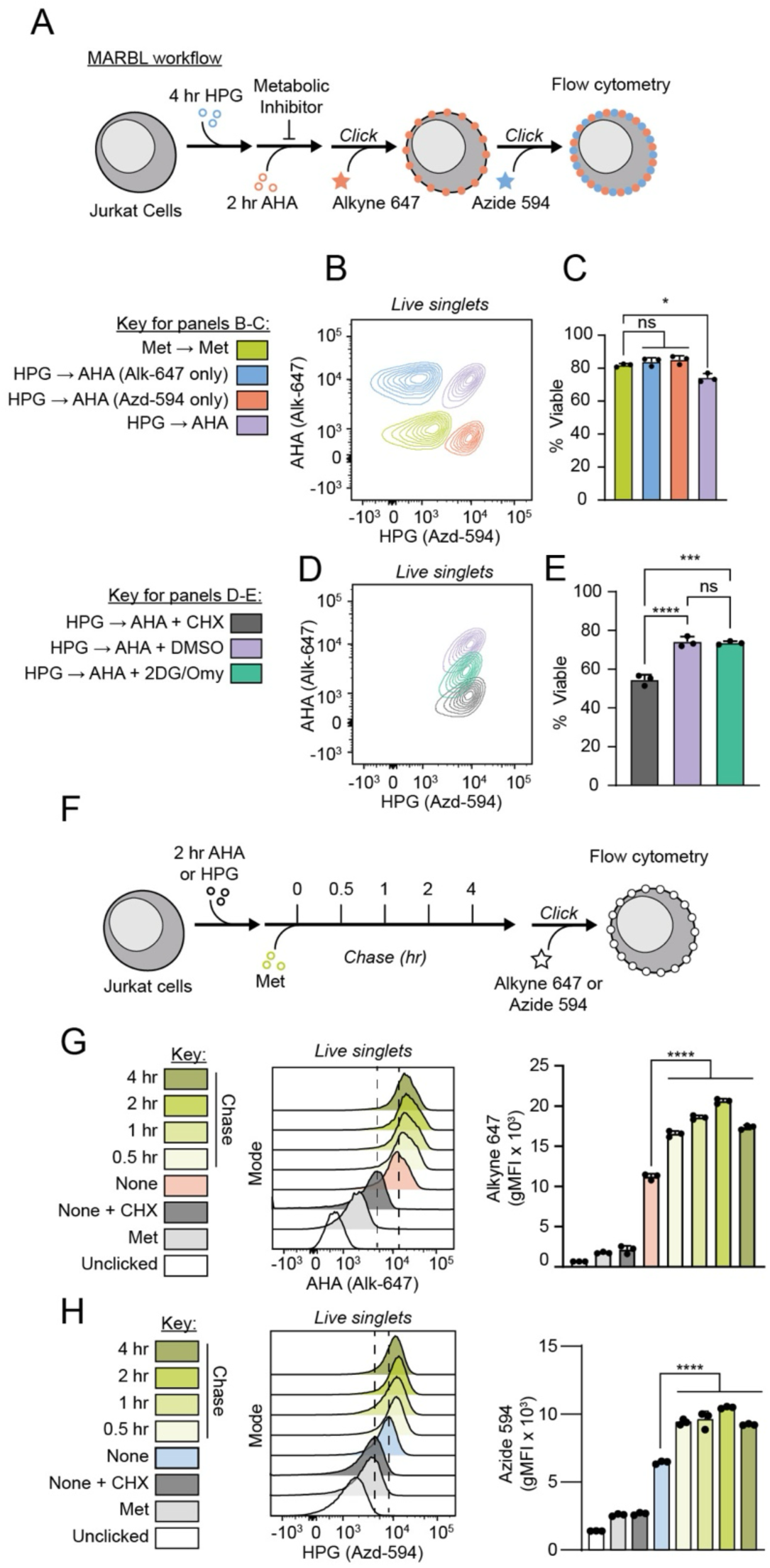
Development of dual-incubation MARBL workflow to measure cellular energetics in single cells. (A) Schematic depicting experimental workflow. (B-C) Representative flow cytometric contour plots of extracellular Alk-647 and Azd-594 signal on Jurkat cells (B) and corresponding cell viability after click staining (C). (D-E) Representative flow cytometric contour plots of extracellular Alk-647 and Azd-594 signal on Jurkat cells (D) and corresponding cell viability after click staining (E) in the presence of vehicle, translation (CHX = 500 µM), or metabolic (2DG = 25 mM, Omy = 2 µM) perturbation. (F) Schematic depicting experimental workflow. (G) Representative flow cytometric histograms and corresponding gMFI of extracellular Alk-647 signal with an AHA pulse followed by a Met chase. (H) Representative flow cytometric histograms and corresponding gMFI of extracellular Azd-594 signal with an HPG pulse followed by a Met chase. Abbreviations: AHA = Azidohomoalanine, HPG = Homopropargylglycine, Met = Methionine, CHX = Cycloheximide, gMFI = geometric mean fluorescence intensity, 2DG = 2-Deoxy-D-Glucose, Omy = Oligomycin A Statistical significance was assessed by one-way ANOVA (C, E, G-H) followed by Tukey’s multiple comparisons test. Graphs display mean ± SD (C, E, G-H). (ns, p >0.05; *, p ≤ 0.05; **, p ≤ 0.01; ***, p ≤ 0.001; or ****, p ≤ 0.0001).

For accurate assessment of translation rates, the CMA signal must remain stable for the duration of the experiment. To assess the stability of CMAs in surface proteins, Jurkat cells were pulsed with AHA or HPG followed by a chase with methionine-supplemented media before performing the click reaction (Fig. 3F). The AHA and HPG signals continued to increase for 0.5 hours during the methionine chase (Fig. 3G-H). We speculate this increase in signal is caused by CMAs incorporated into proteins that have yet to be trafficked to the cell surface. The signal remains stable for 2 hours thereafter and then begins to wane to a small degree between two and four hours of methionine chase (Fig. 3G-H). However, the CMA signal remains baseline-resolved after four hours in methionine. Thus, the CMA signal is durable over the course of the dual-labeling experiment.

### Application of MARBL to primary human immune cells

We next sought to calculate a normalized relationship between baseline and metabolically coupled translation measurements as a single metabolic resilience index per cell (Fig. S4A). In this calculation, raw AHA and HPG signals are variance stabilized by a Yeo-Johnson transformation before filtering out the top and bottom 2.5% outliers in signal (Fig. S4B-S4C). Next, we calculate the difference between the HPG and AHA signals, yielding a single resilience index per cell. Mapping the metabolic resilience index onto a scatterplot of AHA versus HPG signal reveals how this value decreases in response to energetic inhibition (Figs. S4D-S4E). There is an overall decrease in resilience index during Omy and 2DG treatment (Fig. S4F). Comparing the resilience index across an array of drug treatments, reveals that Jurkat cells are insensitive to most single-agent energetic inhibitors (Fig. S4F). This reflects the ability to maintain high translation rates during drug treatment, which may indicate adaptation to the metabolic stress or low engagement of the pathway. Here, we provide an analytical framework to integrate baseline HPG and metabolically coupled AHA incorporation to measure metabolic resilience.

Next, we applied MARBL to a mixed population of primary human cells, in which we could combine an immunophenotyping flow cytometry panel to associate CMA signal with cell type and activation state. Human peripheral blood mononuclear cells (PBMCs) were stimulated and processed with MARBL (Fig. 4A). The low AHA gate captures low-translating cells and overlaps with background signal, whereas the high AHA gate comprises actively translating cells in our vehicle control (Fig. 4B-C). High cell viability was maintained after dual-labeling and extracellular clicking (Fig. S4G). Considering the AHA signal, ∼60% of CD3^+^ T cells fall into the low translation bin and ∼40% are in the high translation bin without metabolic inhibition (Fig. 4C). As expected, addition of CHX halts incorporation of AHA (Fig. 4D). The addition of metabolic perturbations reveals that CD3^+^ T cells are highly sensitive to the inhibition of OXPHOS and largely insensitive to inhibition of glucose metabolism (Fig. 4E-F). Next, we stratified CD3^+^ T cells by the expression of CD69, which is a surface marker for activation, and evaluated the relative representation of T cells within the low versus high AHA gate. Within CD3^+^ T cells, approximately 70% are CD69^-^and 30% are CD69^+^ (Fig. 4G). CD69^-^ cells are enriched in the low AHA bin whereas CD69^+^ cells are enriched in the high AHA bin independent of metabolic perturbation (Fig. 4G). Nevertheless, both CD69^+^ T cells and CD69^-^ T cells display similar metabolic resilience index to Omy and 2DG treatment (Fig. 4H, S4H). These data highlight the importance of ratiometric analysis for single cell interpretation. Without considering baseline translation, we would describe CD69^-^ T cells as inherently vulnerable to Omy treatment, while this instead reflects that CD69^-^ cells are less translationally active.

**Figure 4:**
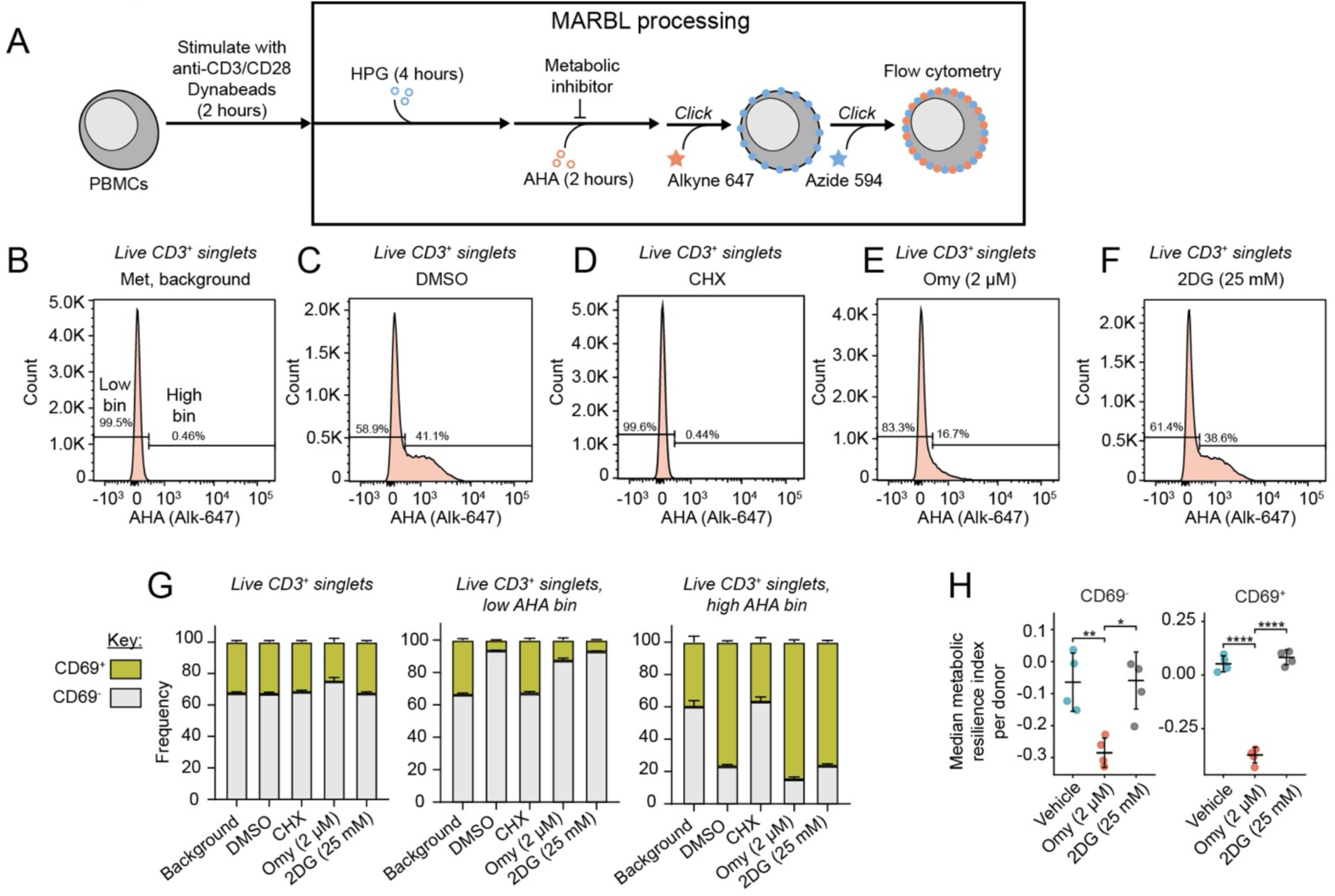
Application of MARBL to primary human cells. (A) Schematic depicting experimental workflow. (B-F) Representative flow cytometric histograms of extracellular Alk-647 signal on activated CD3^+^ human T cells incubated with the following: Met (B), AHA + DMSO (C), AHA + CHX (D), AHA + Omy (E), or AHA + 2DG (F). (G) Quantification of CD69^+^ and CD69^-^ frequency in CD3^+^ singlet gate (left), low AHA bin within CD3^+^ singlet gate (middle), and high AHA bin within CD3^+^ singlet gate (right). (H) Metabolic resilience indices for human CD3^+^ T cells stratified by CD69 activation status. Each dot depicts a different healthy human donor. Experiments (B-H) were performed in biological quadruplicate (n = 4). (I) Abbreviations: CHX = Cycloheximide, 2DG = 2-Deoxy-D-Glucose, Omy = Oligomycin A, gMFI = geometric mean fluorescence intensity, PBMC = Peripheral blood mononuclear cells Statistical significance was assessed by one-way ANOVA (H) followed by Tukey’s multiple comparisons test. Graphs display mean ± SD (G-H). (ns, p >0.05; *, p ≤ 0.05; **, p ≤ 0.01; ***, p ≤ 0.001; or ****, p ≤ 0.0001).

### MARBL reveals energetic dependencies that stratify cell function

A key advantage with MARBL is that it returns live cells for subsequent analysis, such that energetic phenotypes can be linked to cellular functions. To demonstrate this, we first differentiated murine TH17 cells under non-pathogenic (npTH17) or pathogenic (pTH17) polarizing conditions to generate populations of immune cells of the same lineage but distinct functional states. Confirming successful differentiation, pTH17 cells expressed T-bet (Fig. S5A) and both expressed RORγt (Fig. S5B) (33,34). We then processed each cell population with MARBL under vehicle or 2DG/Omy treatment, stained each sample with a CD45 antibody conjugated to different fluorophores, and then mixed npTH17 and pTH17 cells together in equal proportions (Fig. 5A). This mixed population was then sorted by metabolic resilience using Fluorescence Activated Cell Sorting (FACS), using an “all-live” gate to capture the combined population and a “resilient” gate to enrich for cells that maintained translation despite 2DG/Omy treatment (Fig. 5B, S5C). Cells maintained high viability even after clicking and sorting (∼75-85%) (Fig. S5D). While there is little difference in the ratio of pTH17 to npTH17 cells in the “resilient” gate compared to “all-live” gate for the vehicle control, there is ∼four-fold enrichment of pTH17 cells with 2DG/Omy treatment (Fig. 5C, S5E). After resting the sorted cells overnight, they were re-stimulated with PMA/ionomycin to assess the capacity for cytokine secretion. Cells processed with MARBL remained sufficiently viable for this downstream analysis, although viability was reduced relative to unclicked controls (60% to 40%) (Fig. S5F). Supernatants from “resilient” cells contained more interferon-gamma (IFNγ) after re-stimulation, consistent with the reduction of npTH17 and enrichment of pTH17 cells in this gate (Fig. 5D) (35–39).

**Figure 5:**
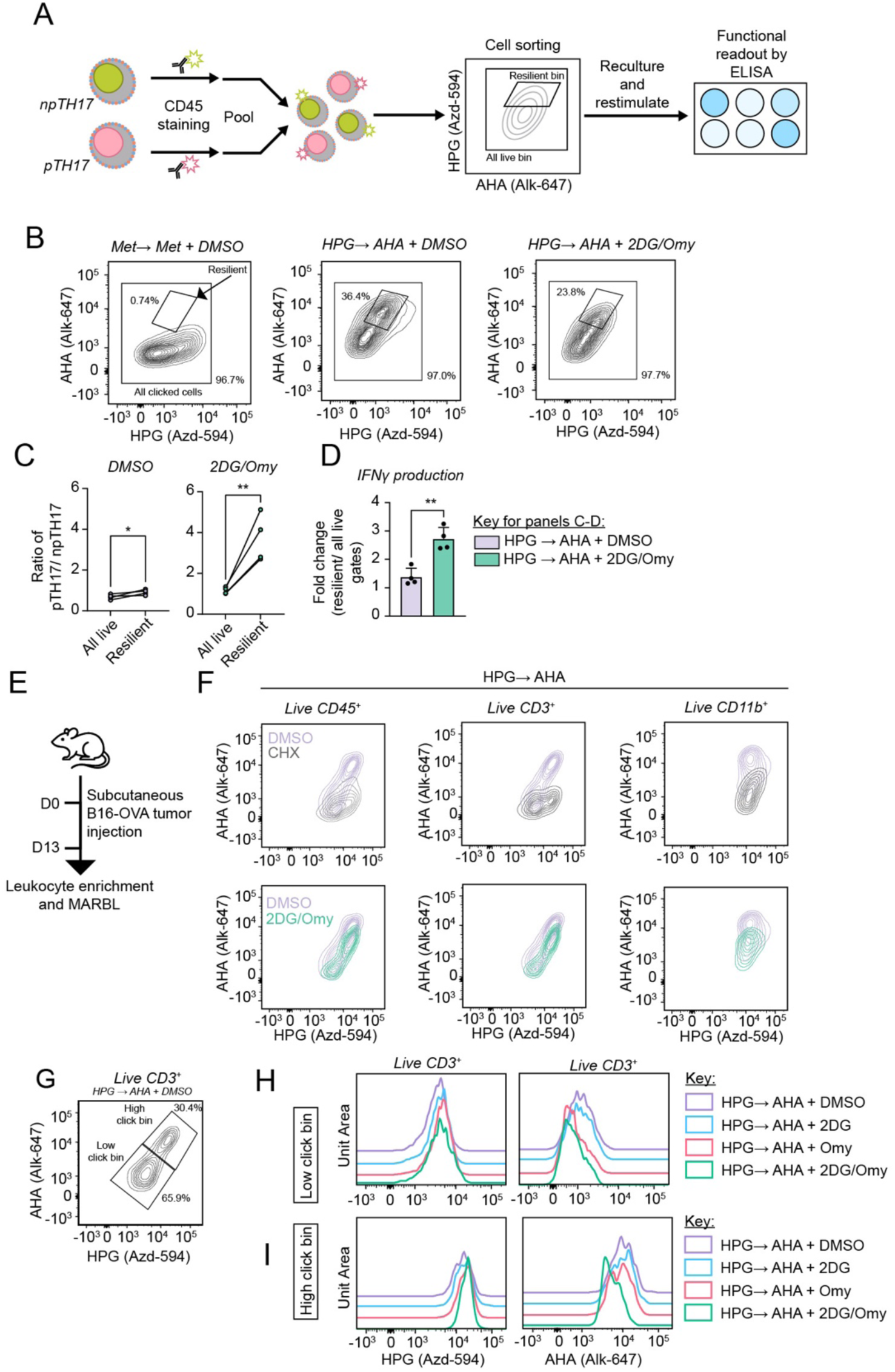
MARBL reveals energetic dependencies that stratify cell function. (A) Schematic depicting experimental workflow. (B) Representative contour plots depicting gates for cell sorting based on extracellular Azd-594 and Alk-647 signals on a 1:1 mixture of npTH17 and pTH17 cells. (C) Ratio of pTH17 cell counts relative to npTH17 cell counts in all live bin and resilient bin. (D) Fold change of IFNγ production following PMA/Ionomycin restimulation comparing resilient gate to all live bin. (E) Schematic depicting experimental workflow. (F) Representative flow cytometric histograms of MARBL-processed leukocytes enriched from B16-Ova tumors in the presence of vehicle treatment, translation inhibition (CHX = 500 µM), or metabolic perturbation (2DG = 25 mM, Omy = 2 µM). (G) Gating strategy for low and high click bins for MARBL-processed CD3^+^ lymphocytes with extracellular Azd-594 and Alk-647 signal. (H-I) Representative flow cytometric histograms of extracellular Azd-594 and Alk-647 signal in low (H) and high (I) click bin gates in the presence of vehicle treatment, translation inhibition (CHX = 500 µM), or metabolic perturbation (2DG = 12.5 mM and Omy = 1 µM). Abbreviations: npTH17 = non-pathogenic TH17, pTH17 = pathogenic TH17, AHA = Azidohomoalanine, Met = Methionine, CHX = Cycloheximide, 2DG = 2-Deoxy-D-Glucose, Omy = Oligomycin A Statistical significance was assessed by two-tailed Student’s unpaired t-test (C-D). Graphs display mean ± SD (C-D). (ns, p >0.05; *, p ≤ 0.05; **, p ≤ 0.01; ***, p ≤ 0.001; or ****, p ≤ 0.0001).

Next, we evaluated the performance of MARBL on an inherently complex biological mixture like the tumor microenvironment. We subcutaneously injected B16-OVA cancer cells into mice, which is a syngeneic melanoma cell line expressing a strong model T cell antigen, performing MARBL on tumor-infiltrating leukocytes ∼2 weeks post-implantation (Fig. 5E). We observed variation in baseline translation among different immune cell populations (Fig. 5F). There is improved incorporation of CMAs into lymphocytes compared to myeloid cells. Next, we evaluated how metabolic responsiveness differed for CD3^+^ T cells with low or high baseline translation (Fig. 5G). Both populations were sensitive to combined 2DG/Omy treatment (Fig. 5H-I). However, only T cells with low translation were sensitive to singular inhibition of OXPHOS with Omy (Fig. 5H). Across all perturbations, baseline HPG signal remained stable (Fig. 5H-I). Together, these findings demonstrate that MARBL enables the recovery of metabolically distinct cell populations from heterogeneous mixtures that remain viable and functional.

## Discussion

MARBL is a flexible, stable, and internally normalized flow cytometry-based platform that combines CMA incorporation with classical perturbational bioenergetics to monitor metabolic dependencies approaching single-cell resolution. By leveraging the unique chemistries of HPG and AHA, we can encode baseline and metabolically coupled incorporation simultaneously into the cell surface proteome. MARBL yields viable cells that can be re-cultured and integrated with other workflows requiring live cells to achieve layered insight. Thus, MARBL expands the toolkit for interrogating metabolic heterogeneity within complex cell populations and can be further multiplexed with downstream assays to identify functional consequences linked to bioenergetic adaptations.

Our method has been extensively optimized to maximize signal over noise and benchmarked to ensure our readout accurately reflects bioenergetic state. In refining this method, we identified culture conditions that enable CMA incorporation to be detected on the cell surface within a few hours and optimized copper-catalyzed click chemistry conditions that are minimally toxic to mammalian cells. CMA incorporation under different metabolic perturbation conditions tracks with the ECAR readout from the Seahorse glycolysis stress test and the adenylate energy charge (AEC) as detected by LC-MS/MS. Furthermore, we successfully applied MARBL to a diversity of sample types, ranging from immortalized cell lines to primary murine or human cells. The breadth and levels of baseline translation varied between cell types, highlighting the importance of our dual-labeling methodology to uncover true metabolic adaptations. The addition of immunophenotyping markers into a flow cytometry panel enables MARBL to be applied to mixed cell populations, stratifying click signal by lineage or activation parameters. Moreover, we sorted, re-cultured, and assessed cytokine production on MARBL-processed npTH17 and pTH17 cells, revealing that metabolic resilience identifies functionally distinct cellular states. Together, these advances establish MARBL as a robust platform for resolving metabolic heterogeneity and coupling bioenergetic state to cellular function.

MARBL builds upon a growing suite of single-cell technologies for profiling metabolism and bioenergetics. Clickable choline incorporation has enabled organelle-selective labeling of phosphatidylcholine (PC), providing a tool to monitor PC transport and trafficking (40,41). Antibody-based approaches such as Met-Flow and scMEP leverage flow cytometry and mass cytometry, respectively, to quantify the expression of key metabolic enzymes and transporters within individual cells (9,10). SCENITH provided the first demonstration that translation rates can serve as a proxy for cellular energetics (11), and several groups have since expanded upon this conceptual framework (Table 1). For example, Cellular Energetics through Noncanonical Amino acid Tagging (CENCAT) replaces puromycin with either HPG or the clickable threonine analogue β-ethynylserine (βES) to monitor translation rates in the presence of different metabolic perturbations (12,29). However, the amino acid analogue signal is still detected within the intracellular proteome, requiring fixation and permeabilization that is incompatible with downstream applications on living cells. By contrast, our MARBL method generates sufficient click signal on the cell surface for flow cytometry and live cell sorting. Lastly, the use of chemically distinct HPG and AHA CMAs in MARBL allows for two conditions to be assessed in the same cell, eliminating the need to split samples into vehicle versus metabolic perturbation conditions and enabling internal normalization and ratiometric analysis. Moreover, the MARBL framework can be readily substituted with other clickable amino acid analogues, such as β-ethynylserine or azidonorleucine, to enable metabolic profiling in native media conditions or an *in vivo* setting, respectively (22,29,42). As a result, MARBL offers an adaptable and complementary strategy for interrogating metabolic dependencies at single-cell resolution in viable populations.

**Table 1:** Comparison of techniques to assay bioenergetics.

| Method | Reporter | Equipment required | Internally controlled | Phenotypic deconvolution | Returns live cells | Compatible with downstream assays | Readout for metabolism | Works in native media |
| --- | --- | --- | --- | --- | --- | --- | --- | --- |
| SCENITH | Puromycin | Flow cytometer | No | Yes | No | Limited | Changes in protein synthesis | Yes |
| Seahorse | ECAR and OCR | Seahorse analyzer | No | No | No | No | Changes in oxygen and pH levels | No |
| CENCAT | HPG or βES | Flow cytometer | No | Yes | No | Limited | Changes in protein synthesis | Depends |
| Met-Flow | Antibody-based detection | Flow cytometer | No | Yes | No | Limited | Changes in protein expression | Yes |
| scMEP | Antibody-based detection | CyTOF mass cytometer | No | No | No | No | Changes in protein expression | Yes |
| MARBL | HPG and AHA | Flow cytometer | Yes | Yes | Yes | Yes | Changes in protein synthesis | No |

MARBL is intended to be a flexible toolkit that can be integrated with any state-of-the-art technique where coupling bioenergetic state with other functional parameters is of interest. For example, future studies may link metabolic phenotypes to genotypes by combining MARBL with genetic perturbation screens, using fluorescently activated cell sorting to enrich for cells with metabolic resilience or sensitivity. As a second application, replacing clickable fluorophores with clickable DNA barcodes may enable MARBL to be applied like CITE-seq (43), adding translationally coupled energetics as another informatic layer in single-cell sequencing datasets. Third, MARBL may be adapted to microscopy platforms to resolve spatiotemporal changes in metabolically coupled protein synthesis (19,21,44). Lastly, the dual-labeling format of MARBL can unmask previously hidden metabolic heterogeneity in cell mixtures of increasing complexity.

A defining feature of MARBL is the resilience index, a new quantifiable characteristic of metabolic responsiveness. The resilience index is independent of the amino acid analogue used to measure translation rates. Consequently, MARBL is a modular technique that can be substituted with other amino acid analogues or click methodology that best suit the question or model system at hand. We anticipate MARBL will enable fresh insight into the metabolic heterogeneity present in complex mixtures in basal and pathological contexts, revealing new mechanisms of cellular adaptation and identifying potential therapeutic vulnerabilities.

## Limitations

Several limitations must be considered when implementing and adapting the MARBL workflow across different sample types. First, efficient incorporation of CMAs requires methionine-free media, as these CMAs exhibit reduced affinity for methionine-tRNA relative to canonical methionine (27). We observed that acute methionine starvation has minimal effects on the polar metabolome at large but does deplete methionine-derived metabolites, which limits the length of incubation before cells adapt to methionine depletion. Second, as MARBL’s readout is the nascent extracellular proteome, CMA incubation times need to be tuned to translational activity to achieve sufficient signal over background, and metabolic perturbations should be optimized to minimize protein turnover. It is important to note that this method may not work for cells with undetectable or low levels of protein synthesis, such as naïve or quiescent cells, and would require prolonged incubation times and/or alternative routes of detection (22,45,46). Moreover, surface proteins have heterogeneous half-lives that can differ across cell types (47). Therefore, absolute gMFI comparisons between cell types should be interpreted with care, as the surface CMA pool integrates over a turnover weighted window that varies by cell type. However, the metabolic resilience index provides a more robust comparison by accounting for cell-type dependent differences in protein turnover. Finally, the current workflow captures primarily acute responses to metabolic perturbation and may require modification of inhibitor dosing or timing. Consequently, highly metabolically plastic cells may evade detection if they can adapt within the duration of our assay.

## Methods

### Experimental Model and Subject Details

#### Cell Lines

Jurkat (ATCC), OCI-AML3, MOLM-13, and K562 cells were cultured in complete RPMI 1640 (10% FBS, 1% penicillin/streptomycin, 10 mM HEPES, 1 mM sodium pyruvate, and 110 μM of β-mercaptoethanol). 143B, Calu6, HCT116, H1299, and B16-Ova cells were cultured in DMEM (Thermo Fisher Scientific, Catalog No.: 11-995-073) supplemented with 10% FBS (R&D Systems, Catalog No.: S11550), and 1% penicillin/streptomycin (Gibco, Catalog No.: 15140-122). All cells were cultured in a humidified 37°C, 5% CO_2_ incubator. Jurkat, OCI-AML3, MOLM-13, HCT116, and H1299 cells are male. K562, 143B, and Calu6 cells are female. All cell lines were free of mycoplasma contamination, assessed by routine PCR-based screening (Bulldog Bio, Catalog No.: 2523448).

#### Mice

Female mice were purchased from Jackson Laboratories (Strain: C57BL/6J, Catalog No. 000664) and housed within the HPPF barrier facility at the Ragon Institute of Mass General, MIT, and Harvard. All mice were female and used between 8-12 weeks of age. All experiments were performed under institutional IACUC approval (MGH protocol: 2022N000056).

### Human Peripheral Blood Mononuclear Cells (PBMCs)

Leukopacs from healthy donors were obtained from the Massachusetts General Hospital Blood Transfusion Service with institutional approval under IRB protocol number: 2005P001218. Prior to blood donation, the donors signed an attestation/consent statement, as per hospital requirements. It states: “I give permission for my blood to be used for transfusion to patients or for research”. All samples are de-identified (completely anonymous), and both the gender and age are not recorded.

### Human PBMC isolation

Leukocytes were diluted 1:1 with 2% heat-inactivated FBS in 1X PBS. Peripheral blood mononuclear cells (PBMCs) were isolated by centrifugation through a Lymphoprep density gradient medium [Stem Cell Technology, Catalog No.: 18060 in SepMate isolation tubes (Stem Cell Technology, Catalog No.: 85450)] at 1200x *g* for 10 minutes. Red blood cells were lysed using ACK lysing buffer (Gibco, Catalog No.: A1049201) for 5 minutes at room temperature.

Isolated PBMCs were resuspended in freezing media (90% FBS, 10% DMSO), placed in CoolCell cell freezing containers (Corning, Catalog No.: 07-210-002), and frozen overnight at −80°C before moving to liquid nitrogen storage.

### Live Cell Labeling with MARBL

Cell Preparation: For adherent cell lines, cells were seeded in 6-well plates one day prior to the labeling experiment at densities that yielded ∼80-90% confluency at the time of labeling. Suspension cells were diluted to a cell density of 1-2×10^6^ live cells/mL and incubated in 96-well V-bottom plates. Cells were washed twice with 1X PBS to remove residual methionine before CMA labeling. For CMA labeling, Jurkat, OCI-AML3, MOLM-13, K562, murine, and human cells were incubated in methionine-free RPMI 1640 (Sigma-Aldrich, Catalog No.: 07-210-002) supplemented with 10% dialyzed FBS (Gibco, Catalog No.: 26400044), 10 mM HEPES (Gibco, Catalog No.: 15-630-080), 1% penicillin/streptomycin (Gibco, Catalog No.: 15140-122), 1 mM sodium pyruvate (Gibco, Catalog No.: 11-360-070), 2 mM glutamine (Gibco, Catalog No.: 25030081), and 200 μM L-Cystine (Alfa Aesar Thermo Fisher Scientific, Catalog No.: A13762.18). Cystine was dissolved in 0.1 M HCl at 50 mM in a heated bath sonicator (Branson) before being diluted to 10 mM with water and filtered through a 0.22 μM polyethersulfone membrane (Nest, Catalog No.: 380111). 143B, H1299, HCT116, and Calu6 cells were incubated in methionine-free DMEM (Thermo Fisher Scientific, Catalog No.: 07-210-002) supplemented with 10% dialyzed FBS, 10 mM HEPES, 1% penicillin/streptomycin, 4 mM glutamine, and 200 μM L-Cystine. Methionine (Sigma-Aldrich, Catalog No.: M9625-5G), azidohomoalanine (Vector Laboratories, Catalog No.: CCT-1066), or homopropargylglycine (Vector Laboratories, Catalog No.: CCT-1067) were added at a final concentration of 100 μM and then incubated in a humidified 37°C, 5% CO_2_ incubator unless stated otherwise.

Single-Color MARBL: Cells were labeled for varying lengths of time (indicated in figures or figure legends) in flasks, 6-well, or 96-well plates in methionine-free RPMI 1640 supplemented with methionine, azidohomoalanine or homopropargylglycine. For conditions containing cycloheximide (Thermo Fisher Scientific, Catalog No.: 357420010), this was added at a final concentration of 500 μM throughout the entire labeling duration. Metabolic inhibitors were added during the last hour of incubation, including 0.25-2 μM oligomycin A (Cayman Chemical, Catalog No.: NC2106728) and/or 6.25-25 mM 2-Deoxy-D-Glucose (MedChem Express, Catalog No.: HY-13966). Adherent cells were washed twice with 1X PBS and dissociated from the plate with enzyme-free dissociation buffer (Gibco, Catalog No.: 13151014) for 5 min at 37 °C. After labeling, cells were washed twice with 1X PBS and incubated in clicking solution for 1 minute at room temperature under constant agitation, protected from light. Clicking solutions were prepared immediately before use and reagents were added in the order listed below. Clicking solution for AHA-labeled cells contained 25 μM Copper (II) sulfate (Sigma-Aldrich, Catalog No.: C1297-100G), 125 μM BTTAA (Vector Laboratories, Catalog No.: CCT-1236), 25 μM alkyne-conjugated fluorophore (Vector Laboratories), and 2.5 mM sodium ascorbate (Sigma-Aldrich, A7631-100G) prepared in 1X PBS. Clicking solution for HPG-labeled cells contained 50 μM Copper (II) sulfate, 250 μM BTTAA, 25 μM azide-conjugated fluorophore, and 2.5 mM sodium ascorbate prepared in 1X PBS. Cells were washed twice with 1X PBS prior to staining with LIVE/DEAD Fixable Dye (Thermo Fisher Scientific) diluted 1:600 in 1X PBS for 15 minutes at 4°C. Finally, cells were fixed in 4% paraformaldehyde (Santa Cruz Biotechnology, Catalog No.: sc-281692) for 15 minutes at room temperature or overnight at 4°C and resuspended in either 1% BSA (w/v) in 1X PBS or MACS buffer (1% heat inactivated-FBS and 2 mM EDTA) before data acquisition by flow cytometry. Glucose dependence was calculated using the following formula:

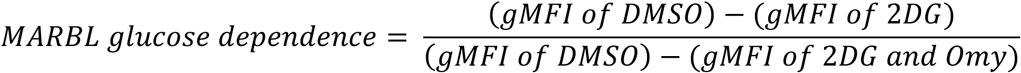

Optimized Dual Color MARBL: Cells were first labeled in HPG-containing culture medium for 4 hours as described above to capture baseline translation. After washing twice with 1X PBS, cells were incubated for 2 hours in AHA-containing culture medium as described above. As described in the previous paragraph, cycloheximide was added at the start of the AHA incubation whereas all metabolic perturbations were added during the final hour. Extracellular clicking reactions were performed sequentially as described above, with two wash steps in 1X PBS in between. Cells were initially clicked with alkyne-conjugated fluorophore followed by azide-conjugated-fluorophore, unless indicated otherwise. Cells were washed twice with 1X PBS prior to staining with LIVE/DEAD Fixable Dye diluted 1:600 in 1X PBS (Thermo Fisher Scientific) for 15 minutes at 4°C. Some samples were stained with extracellular and/or intracellular antibodies. Samples were optionally fixed in 4% paraformaldehyde for 15 minutes at ambient temperature or overnight at 4°C, as described above, or used directly for fluorescence-activated cell sorting. Alternatively, cells were fixed and permeabilized using the Foxp3 Transcription Factor Staining Buffer Set (Thermo Fisher Scientific, Catalog No.: 00-5523-00), according to the manufacturer’s instructions, to enable intracellular antibody staining. Finally, cells were resuspended in MACS buffer before acquisition by flow cytometry.

### Antibody Staining/Flow Cytometry

Live cells were stained with extracellular antibodies for 20-30 minutes at 4°C. The following antibodies were used for extracellular staining: Brilliant Ultra Violet 395 Anti-Mouse CD45.2 (Invitrogen, Catalog No.: 564616), Pacific Blue Anti-Mouse CD45.2 (BioLegend, Catalog No.: 109820), Alexa Fluor® 488 anti-mouse CD45.2 Antibody (BioLegend, Catalog No.: 109815), PE-Cy7 Anti-Mouse CD45.2 (Biolegend, Catalog No.: 109830), Pacific Blue Anti-Mouse/Human CD44 (BioLegend, Catalog No.: 103020), Brilliant Violet 510 Anti-Mouse CD4 (BioLegend, Catalog No.: 100553), Brilliant Ultra Violet™ 496 Anti-Mouse CD11b (Invitrogen, Catalog No.: 364-0112-82), Brilliant Violet 510™ Anti-Mouse CD8b (BioLegend, Catalog No.: 126631), BD Horizon™ Brilliant Ultra Violet™ Anti-Mouse CD3ε (BD Bioscience, Catalog No.: 563565), Brilliant Violet 785™ Anti-Mouse/Human CD44 (BioLegend, Catalog No.: 103059), Brilliant Ultra Violet 395 Anti-Human CD8 (BD Bioscience, Catalog No.: 563795), Brilliant Violet 711 Anti-Human CCR7 (BioLegend, Catalog No.: 353228), APC-Cy7 Anti-Human CD45RA (BioLegend, Catalog No.: 304128), FITC Anti-Human CD4 (BioLegend, Catalog No.: 300538), and PerCP-Cy5.5 Anti-human CD3 (BioLegend, Catalog No.: 344808), which were diluted 1:100 in MACS buffer. For samples with extracellular stain only, cells were fixed in 4% Paraformaldehyde (Santa Cruz Biotechnology, Catalog No.: sc-281692) for 15 minutes at room temperature. In some cases, samples were fixed and permeabilized using the Foxp3 Transcription Factor Staining Buffer Set (Thermo Fisher Scientific, Catalog No.: 00-5521-00) according to the manufacturer’s instructions for intracellular staining. The following antibodies were used for intracellular staining: PE-Cy7 Anti-Mouse Tbet (BioLegend, Catalog No.: 644824), FITC Anti-Mouse FOXP3 (Invitrogen, Catalog No.: 11-5773-83), and PerCP-eFluor 710 Anti-Mouse RORγt (Invitrogen, Catalog No.: 46-6981-82), which were diluted at 1:100 in MACS buffer. Finally, cells were resuspended in MACS (1% HI-FBS, 2 mM EDTA, 1X PBS) buffer before data acquisition by flow cytometry. Precision count beads (BioLegend, Catalog No.: 424902) were optionally added to samples to obtain absolute cell count. Acquisition was performed on the following flow cytometers: BD Fortessa, BD FACS Symphony, or Cytek Aurora. The resulting datasets were analyzed using FlowJo v10.10.0 (FlowJo LLC).

### Confocal Microscopy

Cells labeled with clickable methionine analogues were prepared as described above in single or dual clicking format. All cells were fixed in 4% paraformaldehyde for 15 minutes at room temperature. For intracellular detection, fixed cells were permeabilized in 0.5% Triton X-100 (Sigma Aldrich, Catalog No.: T8787-250ML) diluted in 1X PBS for 15 minutes at room temperature before clicking for 1 minute with a chemically compatible dye. Subsequently, cells were stained with 300 nM of DAPI (Thermo Fisher Scientific, Catalog No.: D1306) diluted in 1X PBS for 5 minutes at 37°C and washed once in 1X PBS. Cells were spun onto poly-L-lysine coverslips (Neuvitro, Catalog No.: NC0818511) at 300 *g* for 3 minutes (0-2 deceleration) and inverted onto slides with Fluoromount G (SouthernBiotech, Catalog No.: OB100-01) mounting media. Images were acquired with a 20X objective and 2X tube lens using the Zeiss Celldiscoverer 7 Microscope through the ZEN microscopy software (ZEISS). Images were processed and scale bars were added with FIJI (open-source software).

### Metabolite Extractions

For polar metabolomics experiments, cells were prepared by resuspending in fresh complete RPMI 1640 medium one day before the experiment. Cells were then switched into methionine-free RPMI 1640 containing no additional amino acids, 100 μM methionine, 100 μM azidohomoalanine, or 100 μM homopropargylglycine as described in the CMA-labeling methods section. Cells were incubated in a humidified 37°C, 5% CO_2_ incubator for the indicated times and then harvested for metabolomics analysis. Cells were pelleted at 400 *g* for 3 minutes at 4°C, washed twice in ice-cold 150 mM ammonium acetate buffer (Sigma-Aldrich, Catalog No.: A7262-500G), and then aspirated of residual wash buffer before storing the resulting pellets at −80°C until metabolite extraction. For ^18^O turnover experiments, 5% (v/v) H_2_^18^O water (Cambridge Isotope Laboratories, Catalog No.: OLM-240-97-1) was added to each sample during the last 5 minutes of incubation to label newly synthesized ATP. A standard curve measuring incorporation at 3- and 6-minute timepoints was prepared alongside the samples.

Cell pellets were resuspended in 200 μL of extraction solvent (containing 4:4:2 [v:v:v:] methanol: acetonitrile: water (containing internal standards). Metabolites were extracted using the freeze-thaw method by freezing cells with liquid nitrogen and thawing in an ice-cold water bath for 5 minutes for three cycles. After, samples were centrifuged at 14,000 RPM for 10 min before the supernatants were transferred to glass LC vials for injection.

### LC-MS Analysis

For widely targeted metabolomics, extracts were analyzed using a Waters Acquity UPLC coupled with AB Sciex 6500+ QTRAP mass spectrometer with a two-column separation method with Multiple Reaction Monitoring (MRM) mode for data collection. First, metabolites were separated with an iHILIC-p column (HILICON). Next, samples were dried and reconstituted in 20% methanol and injected into an ACE® C18-PFP column (Avantor). A pooled quality control (QC) sample was added to the sample list. This QC sample was injected six times for coefficient of variation (CV) calculation for data quality control. Metabolites with CV less than 30% were used for further analysis. These data were analyzed using MultiQuant software (ABSciex) and normalized to the corresponding internal standards.

For ^18^O turnover experiments, extracts were analyzed on an iHILIC-p column (HILICON) on the AB Sciex 6500+ system. To calculate the total AMP, ADP, and ATP levels in the ^18^O-labeled samples, we first calculated the concentration of M0 for AMP (346>79), ADP (426>79), and ATP (506>79) with the external standard curve with 15N5-AMP (MRM 351>79) as the internal standard. Total AMP, ADP, and ATP concentrations were calculated by the ratio of the concentration of M0 and the corresponding enrichment of M0. ^18^O enrichment was calculated by subtracting the signal from unlabeled samples. Adenylate energy charge (AEC) was calculated using the total concentrations of AMP, ADP, and ATP with the following formula:

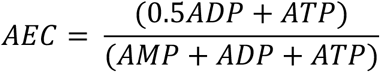

To calculate newly synthesized ATP, we used ATP 239 (3 phosphate tail M0 fragment). The MRMs are 506-239, 508-241, 510-243, and 512-245. Newly synthesized ATP was calculated with the sum of labeled oxygen (enrichment of M2*1+M4*2+M6*3) multiplied by the total ATP concentration. The peak area of each isotopologue was used for the enrichment calculation with the matrix method in accordance with previously published methods (48).

### Cell Growth Curves

2×10^5^ Jurkat cells were washed twice with 1X PBS and incubated in 200 μL of methionine-free media supplemented with methionine, AHA, and/or HPG in 96-well plates. At the indicated time points, cells were mixed 1:1 with a solution containing 5 μg/mL of acridine orange (Thermo Fisher Scientific, Catalog No.: AAAL13159-06) and 100 μg/mL of propidium iodide (Fisher Scientific, Catalog No.: ICN19545825) diluted in 1X PBS. Subsequently, cell viability and number were measured using a CellDrop FL cell counter (DeNovix).

### Mouse tumor model and leukocyte enrichment

Mice were shaved at the injection site and subcutaneously implanted into the abdominal flank with 2.5×10^5^ B16-Ova tumor cells. Once palpable, tumors were monitored and measured with calipers every 2-3 days. Mice were sacrificed day 13 post-injection for tissue harvest. Tumors were suspended in salt-adjusted 2 mg/mL collagenase type I (Worthington, Catalog: NC9482366) dissociated with a gentleMACS™ Dissociator (Miltenyi) using m_imp_tumor_1.1 for 1 minute prior to incubation at 37°C for 20 min. Cells were then processed with m_imp_tumor_3.02 for 37 seconds and filtered through a 70 µM filter. Leukocytes were enriched with a salt-adjusted Percoll (Cytiva, Catalog No.: 17089101) gradient by resuspending dissociated tumor samples in 4 mL of 40% Percoll in 1X PBS and underlaying 2 mL of 70% Percoll in 1X PBS. Samples were centrifuged at room temperature for 20 minutes with the brakes off. Leukocytes were recovered from the hazy interface layer between the 40% and 70% Percoll layers.

### Isolation and stimulation of naïve murine CD4^+^ T cells

Spleens as well as inguinal, axillary, and brachial lymph nodes were harvested from female C57BL/6J mice and mashed through 70 μm nylon mesh filters (Miltenyi, Catalog No.: 130-110-916) to form single cell suspensions. Cells were incubated in ACK lysing buffer for 2 minutes at room temperature to lyse red blood cells. Naïve CD4^+^ T cells were isolated by negative selection according to the manufacturer’s instructions (Miltenyi, Catalog No.: 130-104-453). Naïve CD4^+^ T cells were resuspended in no-glucose RPMI 1640 medium (Gibco, Catalog No.: 11879020) supplemented with 10 mM glucose (Gibco, Catalog No.: A2494001), 10% FBS, 1% penicillin/streptomycin, 10 mM HEPES, 1 mM sodium pyruvate, and 110 μM of β-mercaptoethanol. Then, 5×10^4^ cells were plated onto 96-well flat bottom plates pre-coated with 1 μg/mL anti-CD3 (Bio X Cell, Catalog No.: BE0001-1) and 1 μg/mL anti-CD28 (Bio X Cell, Catalog No.: BE0015-1) antibodies for 96 hours. For pTH17 polarization, the culture medium contained 25 ng/mL of murine IL-6 (BioLegend, Catalog No.: 575702), 20 ng/mL of murine IL-1β (BioLegend, Catalog No.: 575104), and 20 ng/mL of IL-23 (BioLegend, Catalog No. 589002). For npTH17 polarization, the culture medium contained 10 ng/mL of recombinant human TGF-β (PeproTech, Catalog No.: 100-21C) and 25 ng/mL of murine IL-6.

### Human T cell activation

Prior to stimulation, PBMCs were thawed at 37°C and rested overnight in complete RPMI-1640 supplemented with 10% heat-inactivated FBS, 25 mM HEPES, and 1% penicillin-streptomycin. Cells were plated at a density of 1×10^6^ cells per well in a 96-well plate. For PMA/ionomycin stimulation, 50 ng/mL PMA (Sigma-Aldrich, Catalog No.: P8139-1MG) and 1 μg/mL ionomycin (Sigma-Aldrich, Catalog No.: I0634-1MG) were added to the culture medium for 2 hours at 37°C. For polyclonal stimulation, 7.5 μL/well human-activator CD3/CD28 Dynabeads (Gibco, Catalog No.: 11131D) was added to each well for 2 hours at 37°C prior to baseline and metabolically coupled CMA labeling.

### Seahorse Analysis

Oxygen consumption rate (OCR) and extracellular acidification (ECAR) were measured with the Seahorse XF Pro Analyzer (Agilent Technologies, Model No.: S7850A) and Seahorse FluxPaks (Agilent Technologies, Catalog No.: 103793-100). The day prior to metabolic flux analysis, the sensor cartridge was hydrated according to the manufacturer’s instructions. The cell microplate was coated with 50 μL of 50 μg/mL Poly-D-Lysine (Millipore Sigma, Catalog No.: A-003-E) for 1-2 hours at room temperature and washed with 200 μL of water per well. 2×10^5^ total Jurkat cells were plated in 180 μL of XF RPMI medium (Agilent Technologies, Catalog No.: 103576-100) containing 2 mM XF glutamine (Agilent Technologies, Catalog No.: 10-357-9100) in a 96-well cell culture microplate and monitored for 18 minutes (three measurement cycles) under basal conditions and following each injection. Glucose (final concentration of 10 mM), Omy (final concentrations ranging from 0.5-2 μM), and 2DG (final concentrations ranging from 6.25-25 mM) were sequentially injected. Glucose dependence was calculated using the following formula:

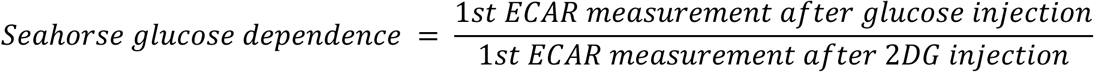

### ELISA Analysis

IFNγ production was measured in culture media using ELISA MAX™ Standard Set Mouse IFN-γ kit (BioLegend, Catalog No.: 430801) according to the manufacturer’s instructions. Data was normalized to absolute live cell counts using Precision Count Beads™ (BioLegend, Catalog No: 424902).

### Cell sorting

Cells were sorted into complete FBS and incubated in complete R10 supplemented with 12.5 ng/mL of IL-2 (Miltenyi, Catalog No.: 130-120-330) overnight. Cells were restimulated with 10 ng/mL of Phorbol 12-myristate 13-acetate (Sigma-Aldrich, Catalog No.: P8139-1MG) and 500 ng/mL of Ionomycin (Sigma-Aldrich, I0634-1MG) for 3 hours. Cell sorting was performed on a BD FACSAria™ Fusion Flow Cytometer. At least 1×10^5^ events were collected for each sort gate.

### Quantification and Statistical Analyses

#### Statistical analysis

Statistics were calculated using GraphPad Prism software (GraphPad Software, Inc., v10). Unpaired Student’s t-test was performed for comparisons between two groups. One-way ANOVA with Tukey’s post-hoc correction was performed for comparisons between three or more groups. Two-way ANOVA with Tukey’s multiple comparisons post-hoc correction was used to compare the impact of two independent variables on a dependent parameter. Bar graphs display mean values, and error bars correspond to standard deviation. The p value notation is as follows: not significant or p >0.05 (ns), * or p ≤ 0.05, ** or p ≤ 0.01, *** or p ≤ 0.001, and **** or p ≤ 0.0001. Heatmaps were created in R (version 4.4.1) using pheatmap (version 1.0.13). For experiments in cell lines, technical replicates and representative results are shown. For experiments in primary cells, biological replicates are shown.

### Calculating Methionine Content in Extracellular vs. Intracellular Domains

Proteome data for humans (UP000005640) and mice (UP000000589) was downloaded from UniProt and filtered for reviewed entries. The human proteome was merged with the membraneome dataset from ETH Zürich’s Wollscheid Lab (https://wlab.ethz.ch/surfaceome/). The mouse proteome was filtered by whether there was a defined topological domain in the UniProt data. Extracellular and intracellular boundaries were determined and used to calculate the amino acid content of the relevant regions in each protein.

### Metabolic Resilience Index

All analyses were performed in R (v4.4.1) within RStudio (v2024.4.2.764) using the bestNormalize (v1.9.1), dplyr (v1.1.4), ggplot2 (v4.0.0), scales (v1.4.0), and scico (v1.5.0.9000) packages. Flow cytometry datasets were exported as CSV files from FlowJo v10.10.0 (FlowJo LLC). and then loaded into RStudio. DMSO-treated control samples were used as a reference to estimate Yeo-Johnson transformation parameters for the AHA and HPG signals separately (using bestNormalize; ε = 0.001; standardize=TRUE), which were then applied to all samples for variance stabilization. Each sample was then scaled to the standard deviation. To minimize the influence of outliers, values outside the 2.5th-98.5th percentile range within each sample were removed. The resulting datasets were used to calculate the metabolic resilience index as the difference between the Yeo-Johnson transformed and scaled values (AHA-HPG) per cell. Metabolic “resilience” indicates the ability of a cell to maintain high translation rates upon drug treatment.

## Acknowledgements

We are grateful for the resources provided by the Ragon Institute Flow Cytometry Core and the Ragon Institute Microscopy Core. Thank you to Yadira Soto-Feliciano, PhD for gifting OCI-AML3, MOLM-13, and K562 cells and Matthew Vander Heiden, MD, PhD for gifting 143B, Calu6, HCT116, and H1299 cells. This material is based upon work supported by the National Science Foundation Graduate Research Fellowship Program under Grant No. (2141064). Any opinions, findings, and conclusions or recommendations expressed in this material are those of the author(s) and do not necessarily reflect the views of the National Science Foundation. A.E.R. acknowledges funding support from the Elsa U. Pardee Foundation Grant and the National Cancer Institute (P30CA14051). I.J.K and Y.Q. were supported by the Stable Isotope and Metabolomics Core Facility of the Diabetes Research and Training Center (DRTC) of the Albert Einstein College of Medicine (National Institutes of Health, United States; P60DK020541). The ABSciex® 6500+ QTrap Mass Spectrometer used in this study was awarded by NIH Shared Instrumentation (1S10 OD021798-01A1) to I.J.K..

## Data availability statement

Datasets and code reported in this paper are available from the lead contact, A.E.R., upon publication.

## Declaration of interests

There is restriction to the use of MARBL due to a pending patent application to I.J.K, Y.Q., L.R.D., and A.E.R. (U.S. Patent 63/681,574 filed August 9, 2024). All remaining authors declare no competing interests.

---

**Figure S1:**
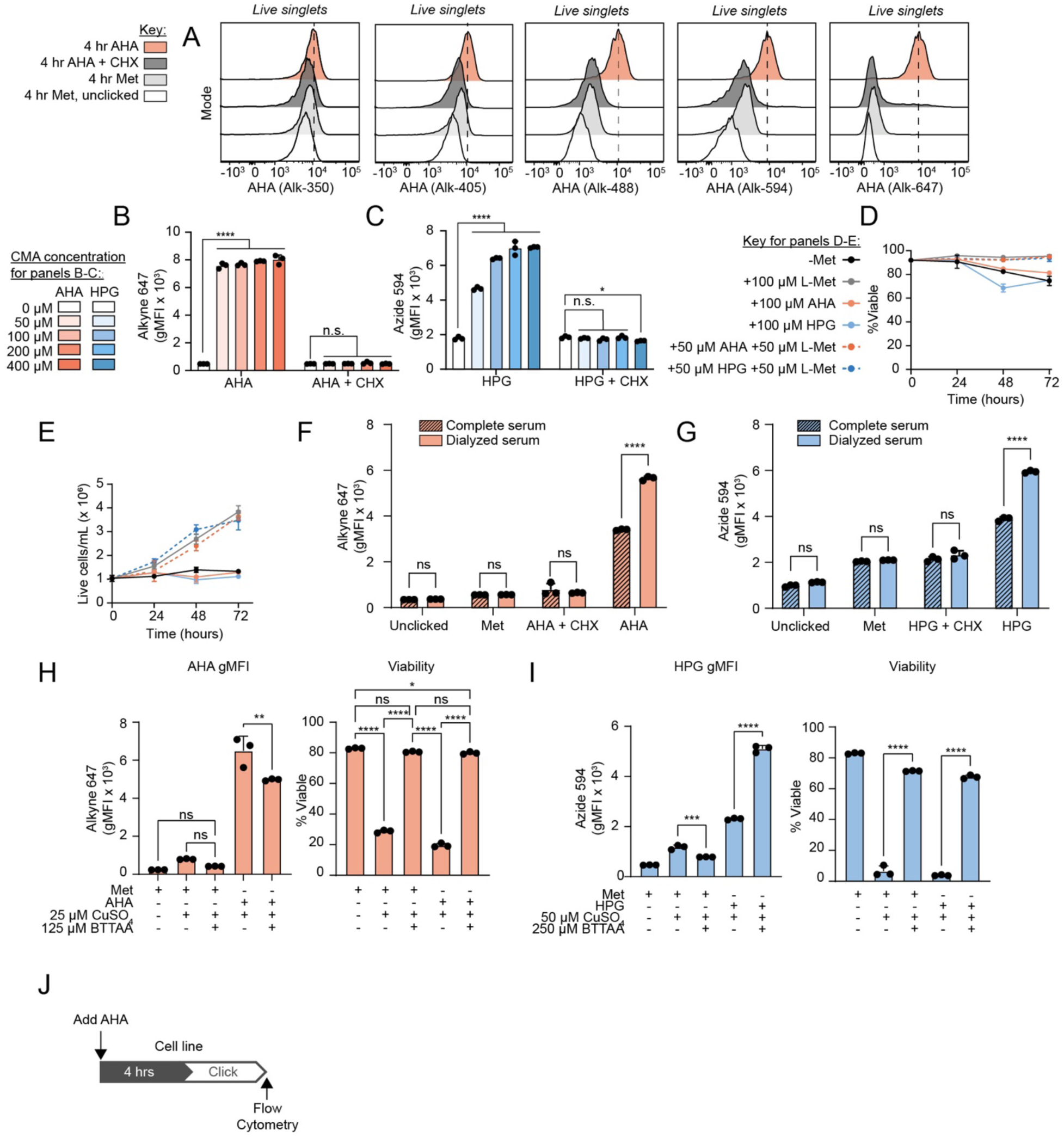
Supporting data for the optimization of clickable methionine analogues to monitor translation in living cells, related to Figure 1. (A) Representative flow cytometric histograms of extracellular Alk-350, Alk-405, Alk-488, Alk-594, or Alk-647 labeled Jurkat cells after 4 hours of AHA incubation. (B) Flow cytometry analysis of Jurkat cells incubated in varying concentrations of AHA or AHA + CHX (500 µM) and extracellularly clicked with Alk-647. (C) Flow cytometry analysis of Jurkat cells incubated in varying concentrations of HPG or HPG + CHX (500 µM) and extracellularly clicked with Azd-594. (D-E) Cell viability (D) and live cell count (E) in Jurkat cells cultured in Met-free media supplemented with Met and/or CMAs. (F-G) Jurkat cells extracellularly clicked with Alk-647 (F) or Azd-594 (G) after 4 hours of CMA incorporation in media supplemented with dialyzed or complete FBS. (H-I) Viability and gMFIs of Jurkat cells incubated in AHA or HPG for 4 hours with varying concentrations of CuSO4-chelating ligand BTTAA and extracellularly-clicked with Alk-647 (H) or Azd-594 (I). (J) Schematic depicting experimental workflow. Statistical significance was assessed by one-way ANOVA (B-C, H-I) or two-way ANOVA (F-G) followed by Tukey’s multiple comparisons test. Confocal images were acquired at 20X magnification with a 2X tube lens (B-C). Graphs display mean ± SD (B-I). (ns, p >0.05; *, p ≤ 0.05; **, p ≤ 0.01; ***, p ≤ 0.001; or ****, p ≤ 0.0001). Abbreviations: AHA = Azidohomoalanine, HPG = Homopropargylglycine, Met = Methionine, CHX = Cycloheximide, gMFI = Geometric mean fluorescence intensity, FBS = fetal bovine serum

**Figure S2:**
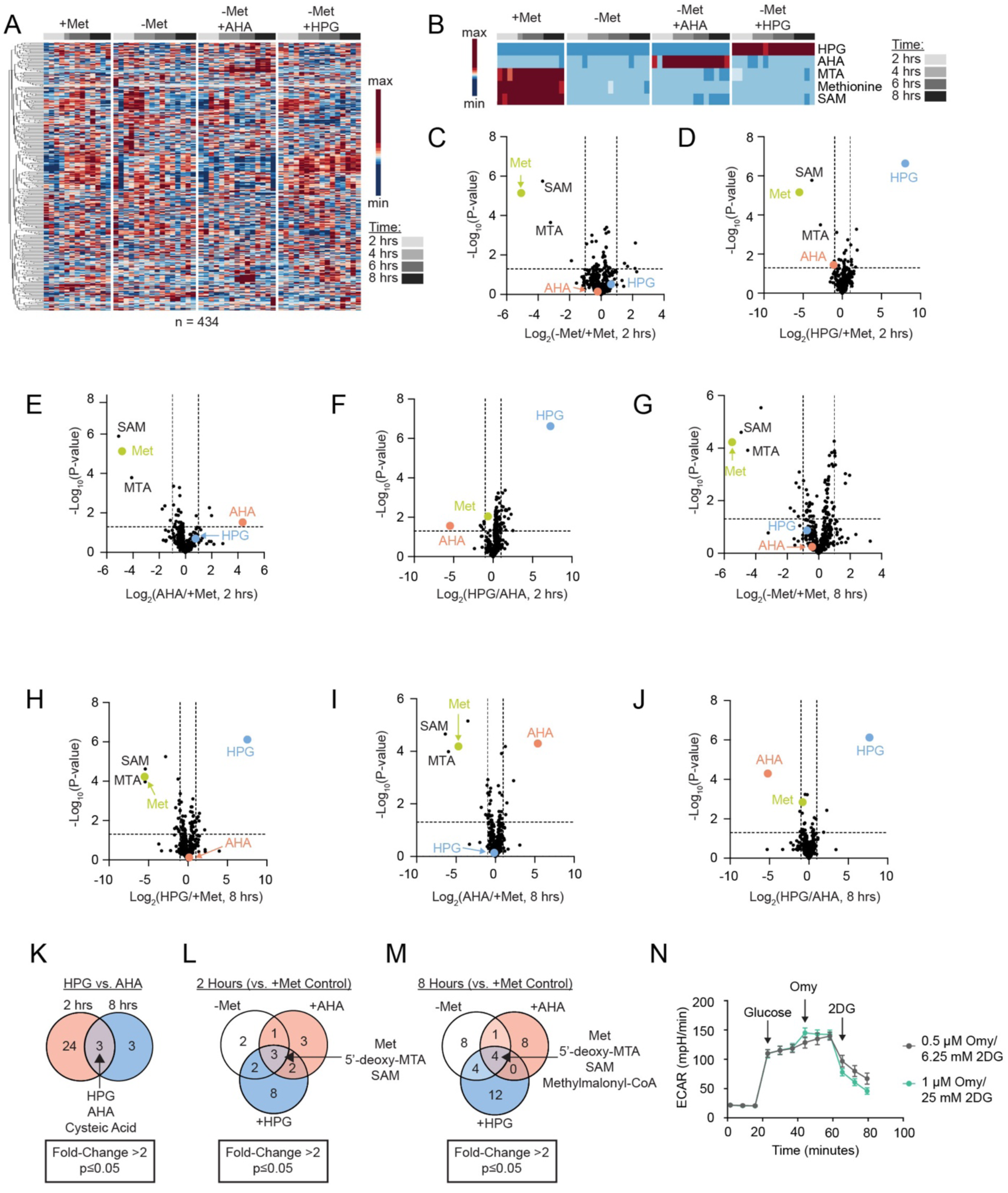
Supporting data for validation of surface translation rate as a readout of cellular energetics, related to Figure 2. (A) Heatmap of polar metabolites after 2, 4, 6, and 8 hours of incubation in Met-free media supplemented with Met, AHA, or HPG. (B) Heatmap of methionine-derived metabolites, AHA, and HPG. (C-J) Volcano plots of metabolites differentially enriched in various media conditions after 2 (C-F) or 8 (G-J) hours of incubation. (K-M) Venn diagrams of significantly altered metabolites in various media conditions using a p-value cutoff of 0.05, comparing: HPG versus AHA after 2- or 8-hour incubations (K); HPG, AHA, or -Met compared to +Met control after a 2-hour incubation (L); and HPG, AHA, or -Met compared to + Met control after an 8-hour incubation (M). (N) Seahorse glycolysis stress test on Jurkat cells using varying concentrations of 2DG and Omy. Abbreviations: AHA = Azidohomoalanine, HPG = Homopropargylglycine, Met = Methionine, 2DG = 2-Deoxy-D-Glucose, Omy = Oligomycin A, MTA = 5’-methylthioadenosine, SAM = S-adenosylmethionine, ECAR = Extracellular acidification rate Statistical significance was calculated using a two-tailed Student’s unpaired t-test (C-M). Graphs display mean ± SD (N). (ns, p >0.05; *, p ≤ 0.05; **, p ≤ 0.01; ***, p ≤ 0.001; or ****, p ≤ 0.0001).

**Figure S3:**
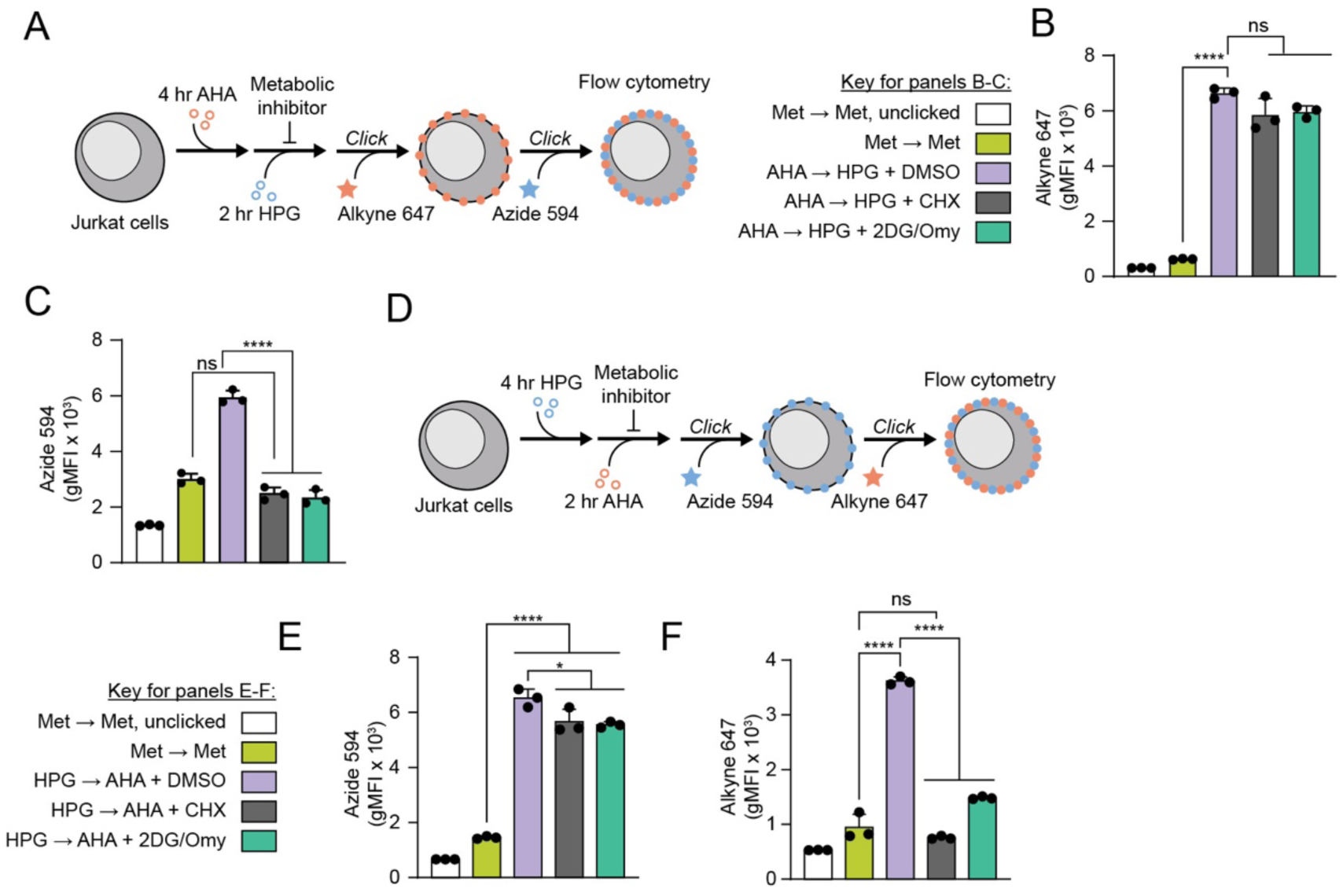
Supporting data for development of dual-incubation MARBL workflow to measure cellular energetics in single cells, related to Figure 3. (A) Schematic depicting experimental workflow. (B-C) Extracellular Alk-647 gMFI (B) and Azd-594 gMFI (C) in the presence of vehicle, translation inhibition (CHX = 500 µM), or metabolic perturbation (2DG = 25 mM, Omy = 2 µM). (D) Schematic depicting experimental workflow. (E-F) Extracellular Azd-594 gMFI (E) and Alk-647 gMFI (F) in the presence of vehicle, translation inhibition (CHX = 500 µM), or metabolic perturbation (2DG = 25 mM, Omy = 2 µM). Abbreviations: AHA = Azidohomoalanine, HPG = Homopropargylglycine, Met = Methionine, CHX = Cycloheximide, gMFI = geometric mean fluorescence intensity, 2DG = 2-Deoxy-D-Glucose, Omy = Oligomycin A Statistical significance was assessed by one-way ANOVA (B-C, E-F) followed by Tukey’s multiple comparisons test. Graphs display mean ± SD (B-C, E-F). (ns, p >0.05; *, p ≤ 0.05; **, p ≤ 0.01; ***, p ≤ 0.001; or ****, p ≤ 0.0001).

**Figure S4:**
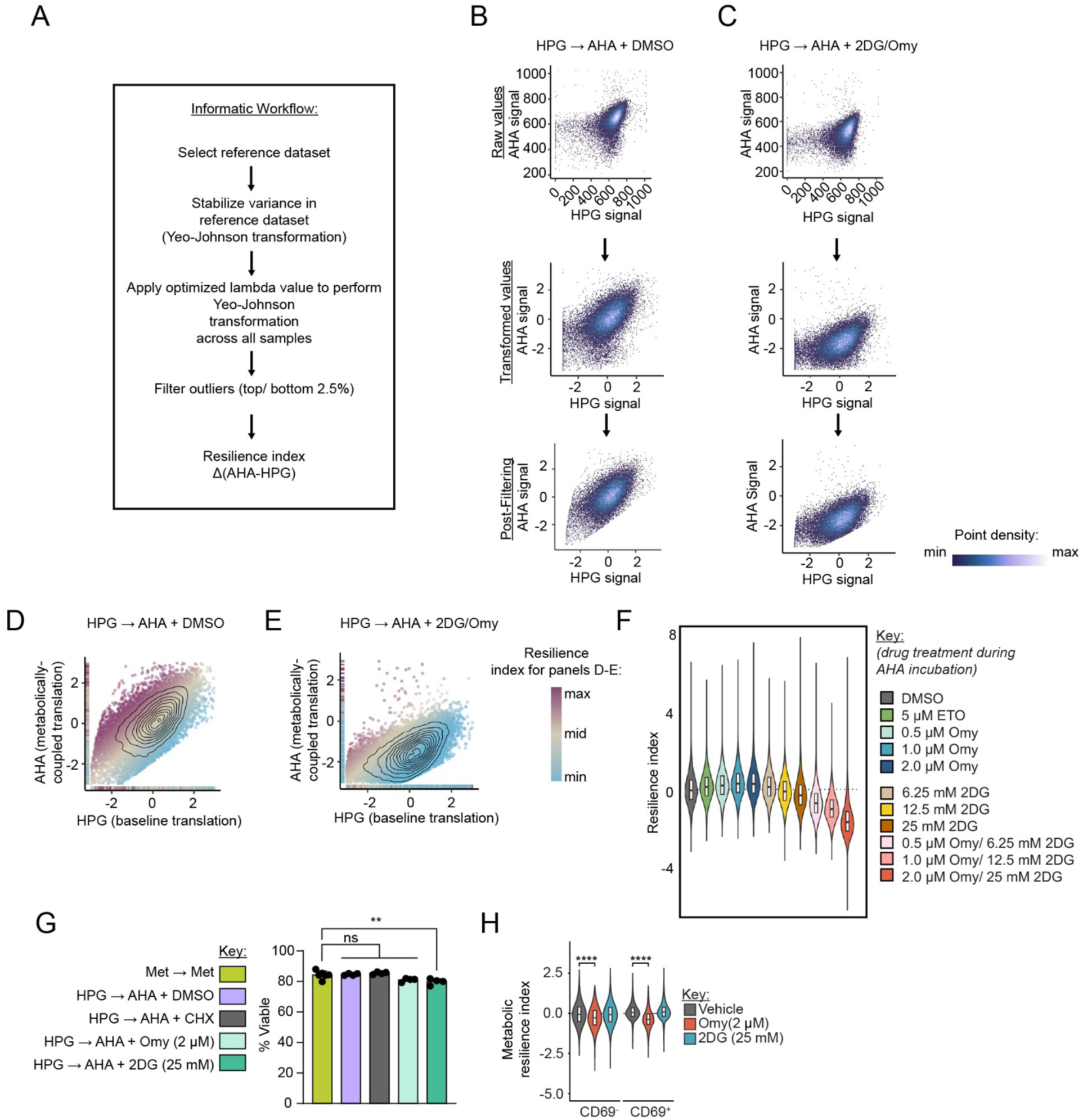
Supporting data for application of MARBL to primary human cells, related to Figure 4. (A) Informatic workflow for calculating metabolic resilience index. (B-C): Transformation and filtering of raw click signal on dually HPG and AHA labeled Jurkat cells treated with vehicle (B) or metabolic inhibitors (2DG = 25 mM, Omy = 2 µM). (D-E) Individual cells colored by metabolic resilience index for cells treated with vehicle (D) or metabolic inhibitors (2DG = 25 mM, Omy = 2 µM) (E). (F) Violin plots of metabolic resilience index of Jurkat cells. (G) Cell viability of PBMCs after dual-labeling and extracellular clicking. (H) Violin plots of metabolic resilience indices of human CD3^+^ T cells stratified by CD69^+^ activation status. Experiments (G-H) were performed in biological quadruplicate with four healthy donors (n = 4). Abbreviations: AHA = Azidohomoalanine, Met = Methionine, CHX = Cycloheximide, 2DG = 2-Deoxy-D-Glucose, Omy = Oligomycin A, ETO = Etomoxir, PBMC = Peripheral blood mononuclear cells Statistical significance was assessed by one-way ANOVA (G-H) followed by Tukey’s multiple comparisons test. Graphs display mean ± SD (G). (ns, p >0.05; *, p ≤ 0.05; **, p ≤ 0.01; ***, p ≤ 0.001; or ****, p ≤ 0.0001).

**Figure S5:**
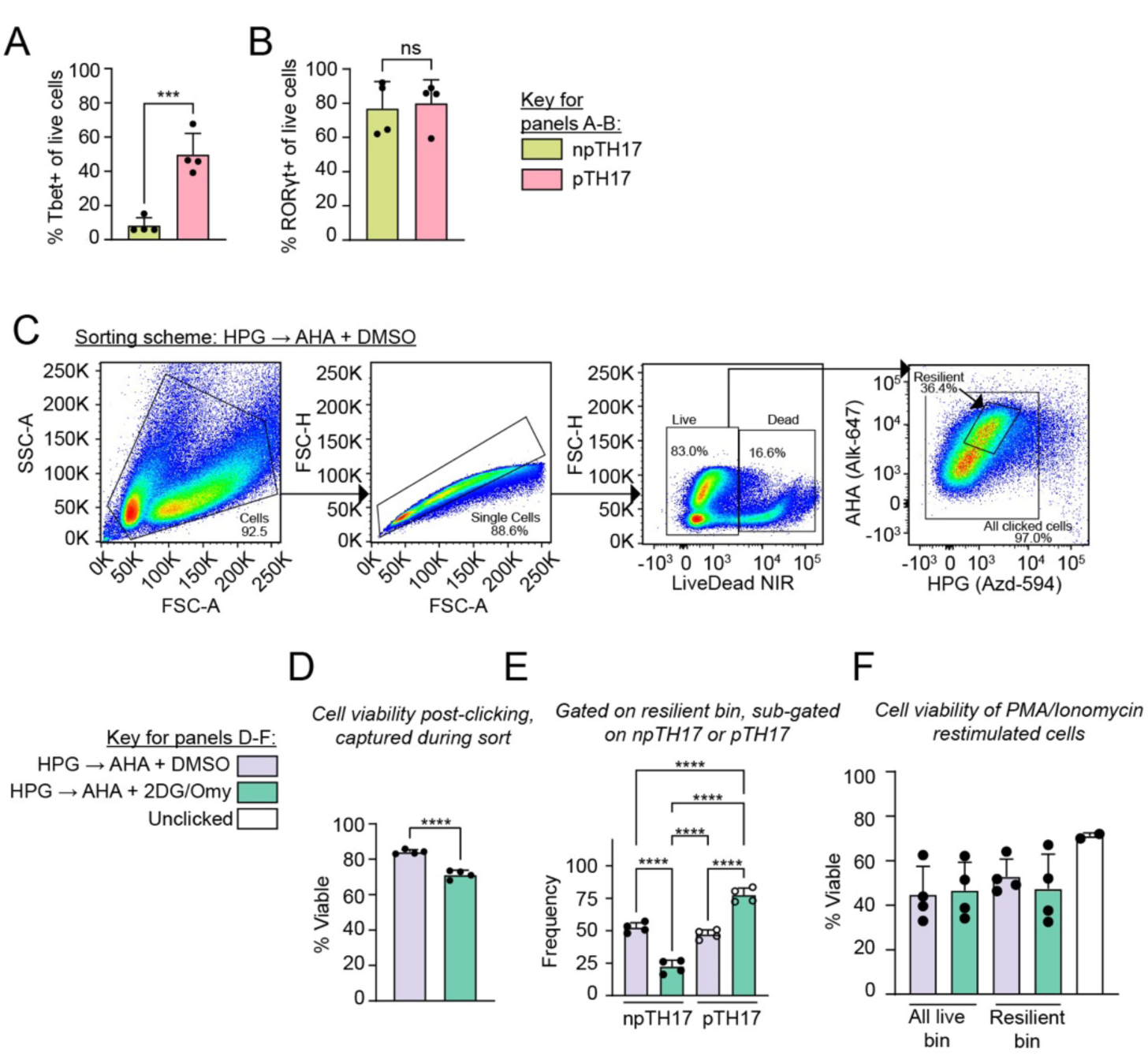
Supporting data for MARBL reveals energetic dependencies that stratify cell function, related to Figure 5. (A-B) Frequency of Tbet^+^ (A) and RORγt^+^ (B) cells in murine naïve, non-pathogenic, or pathogenic CD4^+^ TH17 cells. (C) Complete gating scheme for mixed npTH17 and pTH17 sorting experiment. (D) Viability as measured during cell sorting of MARBL-processed npTH17 and pTH17 cell mixtures. (E) Frequency of dually MARBL-processed npTH17 or pTH17 cells in resilient bin. (F) Viability of MARBL-processed npTH17 and pTH17 cell mixtures following PMA/Ionomycin restimulation. Abbreviations: npTH17 = non-pathogenic TH17, pTH17 = pathogenic TH17, AHA = Azidohomoalanine, Met = Methionine, CHX = Cycloheximide, 2DG = 2-Deoxy-D-Glucose, Omy = Oligomycin A, FSC = Forward scatter, SSC = Side scatter, NIR = Near-IR viability dye Statistical significance was assessed by two-tailed Student’s unpaired t-test (A-B, D) or by one-way ANOVA (E-F) followed by Tukey’s multiple comparisons test. Graphs display mean ± SD (A-B, D-F). (ns, p >0.05; *, p ≤ 0.05; **, p ≤ 0.01; ***, p ≤ 0.001; or ****, p ≤ 0.0001).

